# Plant-invertebrate interactions across a forested retrogressive chronosequence

**DOI:** 10.1101/2022.03.30.486383

**Authors:** Anne Kempel, Eric Allan, Martin M. Gossner, Malte Jochum, David A. Wardle

**Author notes:** Statement of authorship: AK and DW designed the study, AK collected the data, MJ calculated invertebrate biomass, AK analysed the data with substantial input from EA and wrote the manuscript with substantial input from EA, DW, MG and MJ. Data accessibility statement: Should the manuscript be accepted, the data supporting the results will be archived in a public repository (Dryad) and the data DOI will be provided.

## Abstract

In the long-term absence of disturbance, ecosystems often enter a decline or retrogressive phase which leads to reductions in primary productivity, plant biomass, nutrient cycling and foliar quality. However, the consequences of ecosystem retrogression for higher trophic levels such as herbivores and predators, are less clear. Using a post-fire forested island-chronosequence across which retrogression occurs, we show that nutrient availability strongly controls invertebrate herbivore biomass when predators are few, but that there is a switch from bottom-up to top-down control when predators are common. This trophic flip in herbivore control probably arises because invertebrate predators respond to alternative energy channels from the adjacent aquatic matrix, which were independent of plant biomass. Our results suggest that effects of nutrient limitation, following ecosystem retrogression, on trophic cascades are modified by independent variation in predator abundance, which requires a more holistic approach to trophic ecology to better understand herbivore effects on plant communities.

## INTRODUCTION

Nutrient availability and limitation shape ecosystems and food-webs in fundamental ways (Vitousek 2004). Following major disturbances, increases in nutrients initially lead to a build-up in plant biomass and productivity (Peltzer et al., 2010; Wardle et al., 2004). However, in the long-term absence of major disturbance, nutrient limitation increases (Vitousek 2004; Peltzer *et al*. 2010), frequently leading to “ecosystem retrogression” characterized by decreases in plant biomass and primary productivity, and in rates of decomposition and nutrient cycling (Walker & Syers 1976; Wardle *et al*. 2004; Laliberté *et al*. 2013; Turner & Laliberté 2015). While the build-up phase and its consequences for higher trophic levels, such as herbivores and their predators, are relatively well understood (Brown, 1985; Fagan & Bishop, 2000; Neves et al., 2014; Siemann et al., 1999), the consequences of ecosystem retrogression for higher trophic levels are less clear (Gruner 2007; Crutsinger *et al*. 2008). Long-term chronosequences that include retrogressive states can improve our understanding of how herbivore and predator communities are affected by nutrient limitation. This is because they occur over longer time scales and are less impacted by short-term processes typical for the build-up phase such as high turnover in plant species compositions (Peltzer *et al*. 2010). Thus, retrogressive chronosequences can be used as model systems for contributing to the development of general principles about the extrinsic factors that regulate higher trophic levels and their impacts on plants.

As retrogression proceeds and soil fertility declines, plant species composition shifts from species or genotypes with resource-acquisitive traits towards those with more conservative traits (Coley *et al*. 1985; Reich 2014). As such, nutrient concentrations in plant tissues often decrease and plant defences increase (Wardle *et al*. 1997; Hättenschwiler *et al*. 2003; Vitousek 2004). This may influence herbivore preference and performance, and hence herbivore identity, abundance, and impact (Root 1973, Feeny 1976, Kempel et al. 2011). Shifts towards more conservative plant species and reduced plant productivity that accompany declining soil fertility are expected to reduce herbivore biomass (e.g. Chase et al., 2000; McNaughton et al., 1989), and this has frequently been shown in managed grasslands and grassland experiments (Borer *et al*. 2012; Simons *et al*. 2014; Ebeling *et al*. 2018; Welti *et al*. 2020). However, few studies have assessed the relationship between nutrient availability and invertebrate herbivore abundance along natural gradients of nutrient availability (Cuevas-Reyes *et al*. 2004), and the few studies that have been performed across lengthy gradients such as those provided by retrogressive chronosequences have produced mixed results (Gruner 2007; Crutsinger *et al*. 2008).

In addition to bottom-up forces, top-down regulation by predators can strongly affect the biomass of herbivores and their impact on plants (Hairston *et al*. 1960; Barnes *et al*. 2020). Recent findings suggest that the strength of bottom-up control of herbivores depends on predator abundance (Letnic & Ripple 2017; Barnes *et al*. 2020; Welti *et al*. 2020). Similarly, the ‘Exploitation Ecosystems Hypothesis’ (EEH) predicts that bottom-up and top-down forces operate simultaneously, but their relative importance within communities changes with ecosystem productivity (Oksanen *et al*. 1981; Fretwell 1987; Oksanen & Oksanen 2000). In very unproductive systems, predators are predicted to be absent and herbivore biomass therefore increases strongly as plant biomass increases. In more productive systems, herbivore biomass is predicted to show only weak responses to increasing plant biomass as predators control herbivores. This theory was originally formulated for vertebrates, and the “trophic flip” (from bottom-up to top-down) has been shown for vertebrate food webs (e.g. Aunapuu et al., 2008; Crête, 1999; Letnic & Ripple, 2017). For invertebrates, there is some support for the EEH from a study showing that grassland plant biomass only increased grasshopper and Auchenorrhyncha (Hemiptera) biomass when spider biomass was low, but the theory was not fully supported because spider biomass was unrelated to plant biomass (Welti *et al*. 2020). Further, invertebrate predators may not respond strongly to plant biomass if they also feed on detritivores, pollinators or prey from interconnected aquatic systems (Gounand *et al*. 2018), which are not necessarily linked to plant biomass (Clough *et al*. 2014). Hence, whether the trophic flip predicted by the EEH can be found in forested ecosystems, and holds for entire invertebrate communities, remains unexplored.

While few studies have assessed the relationship between herbivore and plant biomass along productivity gradients, our understanding of how the *impact* of herbivores, i.e. their effect on plant biomass or other community components, changes across these gradients is less clear (Coupe & Cahill 2003; Schädler *et al*. 2003; Stein *et al*. 2010; Barnes *et al*. 2020). The resource availability hypothesis (Coley et al. 1985) predicts that impact should be higher in resource rich environments, which select for fast-growing, poorly defended plants (Fine et al. 2004, Blumenthal 2006). However, fast-growing plants typically tolerate herbivores better, as they can compensate for lost biomass (Gianoli & Salgado-Luarte 2017). In contrast, the plant stress hypothesis (White 1969, Rhoades 1983) posits that impact should be higher in more stressful, low-resource environments. Considering that top-down forces increase in parallel with productivity, the EEH states that herbivore impact should show a hump-shaped relationship with productivity (Fraser & Grime 1997; Schädler *et al*. 2003). Several studies use herbivore *damage* to infer *impact* (e.g. Crutsinger et al., 2008; Denno et al., 2002; Ebeling et al., 2021; Endara & Coley, 2011). However, damage by herbivores is not necessarily related to impact because some plants might be more tolerant to enemies than others (Gianoli & Salgado-Luarte, 2017), and the actual *impact* might therefore be lower than expected based on damage alone (Schädler *et al*. 2003). Moreover, the damage caused by sap suckers is difficult to quantify and impact could be greater than expected based on damage. To test for variation in impact, herbivore exclusion studies along soil fertility gradients are necessary, but these have rarely been performed (but see Stein et al., 2010 for grasslands).

In this study, we use a well-studied system of 30 forested lake islands in the boreal forested zone of northern Sweden, for which fire from lightning strikes is the main agent of disturbance. Large islands have burned more frequently than smaller ones due to a higher likelihood of being struck by lightning, resulting in a 5000-year post-fire retrogressive chronosequence (Wardle *et al*. 1997, 2003, 2012). As island size declines, and time since fire increases, nutrient availability declines, and there is a shift to domination by plants with resource-conservative traits and reduced plant biomass and net primary productivity (Wardle *et al*. 2012). On each island, we sampled invertebrates and quantified the biomass of chewing and sucking herbivores, and of predators. In addition, we assessed herbivore damage and impact using herbivore exclusion experiments on phytometer tree seedlings. We also estimated bird predation rate using plasticine caterpillars to test whether herbivore biomass cascades up the food web and is influenced by higher trophic levels. We assessed the biomass of chewing and sucking herbivores separately, as these different feeding guilds may respond differently to variation in plant quality and quantity (Gely *et al*. 2020), and because they may differ in their importance for higher trophic groups (Nyffeler *et al*. 2018). Through structural equation modelling (SEM), we aimed to disentangle the direct and indirect drivers of invertebrate herbivore biomass, damage and impact.

We tested three main hypotheses related to how changes in plant productivity cascade through the food web: (1) Herbivore biomass tracks plant biomass, i.e., herbivore biomass declines with decreasing island size as time since fire increases and nutrients become more limiting. (2) Predator biomass is low on small and unproductive islands, and therefore plant biomass is the dominant control of herbivore biomass, while on large, productive islands predator biomass is higher and suppresses herbivores, resulting in a trophic flip from top-down to bottom up control of herbivores with decreasing island size and increasing nutrient limitation. (3) Herbivore damage decreases with decreasing plant and herbivore biomass as island size decreases (resource availability hypothesis), but herbivore impact does not change because plant tolerance and predator abundance is lower in less fertile environments. In addition, we test a range of other additional hypotheses in our SEM, see Table 1 for all 17 hypotheses that we test. We anticipate that addressing these hypotheses in combination will help advance fundamental understanding of the context-dependency of plant-herbivore-predator interactions.

**Table 1:**
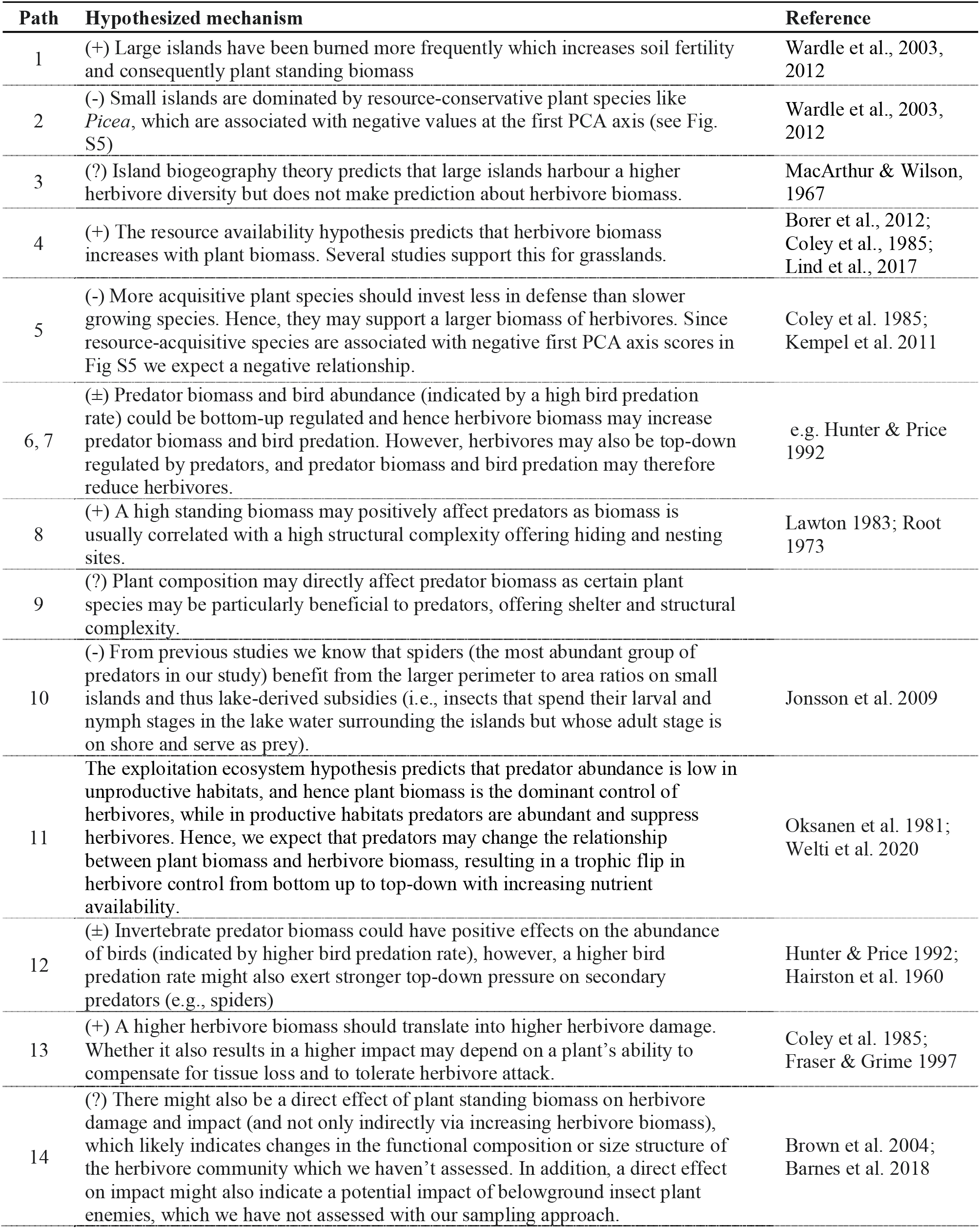
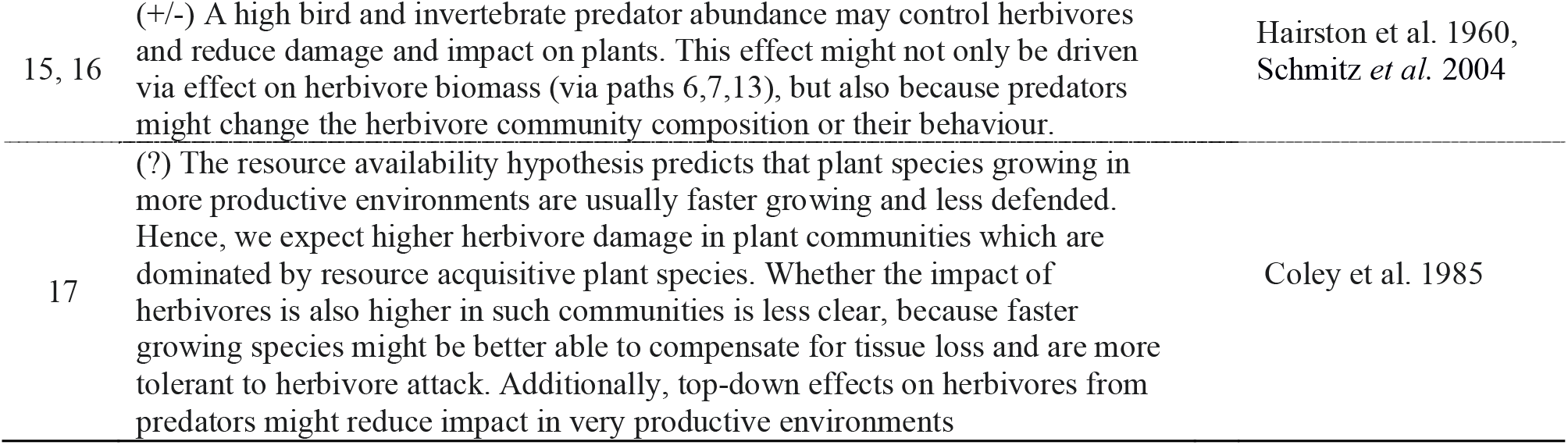
Hypothesized mechanisms affecting herbivore and predator biomass, and herbivore damage and impact, used to underpin the full SEM model that we tested (Fig. 1).

## METHODS

### Island system

We conducted our study on each of 30 forested islands varying in size from 0.03ha to 15ha in two adjacent freshwater lakes in northern Sweden, Lake Hornavan and Lake Uddjaure. The 30 islands have been selected so that their areas are distributed log-normally: ten islands were larger than 1 ha, ten between 0.1 and 1 ha and ten were smaller than 0.1 ha (Wardle *et al*. 2003, 2012, Table S1). All islands were formed following the retreat of land ice about 9000 years ago. The major extrinsic factor that varies among islands is the frequency of fires, which occur as a result of lightning strikes: larger islands are struck more often and have therefore, on average, burned more recently (Wardle *et al*. 1997, 2012). These islands collectively represent a 5000-year chronosequence of increasing time since the most recent fire, and allow insights into important ecological processes over a spatial and temporal scale which would be impossible to assess with experimental approaches (Fukami & Wardle 2005; Walker *et al*. 2010). With increasing time since fire, nutrients become increasingly limiting, which causes a strong decline in plant standing biomass, ecosystem productivity, decomposition rates and nutrient fluxes (Wardle et al., 2012, Table S1).

There are large changes across the island gradient in vegetation composition (Wardle et al. 2012). Specifically, larger, regularly burned islands are dominated by relatively fast-growing plant species with resource-acquisitive functional traits, such as the tree *Pinus sylvestris* and the dwarf shrub *Vaccinium myrtillus* (hereafter *Pinus, V. myrtillus*). Smaller, infrequently burned islands are dominated by slow-growing resource-conservative species, such as the tree *Picea abies* and dwarf shrub *Empetrum hermaphroditum* (hereafter *Picea, Empetrum*). Mid-sized islands are dominated by the tree *Betula pubescens* and dwarf shrub *Vaccinium vitis-idaea* (hereafter *Betula* and *V. vitis-idaea*) which show intermediate growth strategies. Although the relative abundance of the species varies across the island gradient, all species occur on all 30 islands, except for *Pinus* and *Picea* which are absent from five small and three large islands, respectively. Previous work in this system has shown that plants dominating the smaller islands produce foliage of poor quality with high levels of secondary metabolites (Wardle *et al*. 1997; Crutsinger *et al*. 2008). The large changes in soil nutrient availability across the chronosequence, and associated changes in plant biomass and primary productivity, make it an ideal study system for exploring the patterns of plant-herbivore-predator interactions along a natural soil fertility gradient.

### Plant standing biomass and plant functional composition

For each island we obtained data on plant standing biomass per square meter from a previous study in this system which assessed biomass of the six dominant plant species (i.e. *Pinus, Betula, Picea, V. myrtillus, V. vitis-idaea* and *Empetrum*) using allometric equations (Wardle et al., 2003, 2012). To obtain a relative estimate of plant functional composition for each island we ran a principal component analysis (PCA, Fig. S1) across the 30 islands on net primary production (NPP) for these six plant species which collectively account for >98% of total NPP, using data from Wardle et al. (2003). The first PCA axis (PC1) explained 66% of the total variation in the data, and we used PC1 values as an indicator of plant functional composition. High scores of PC1 indicate that the plant community is mostly composed of resource-conservative plant species such as *Picea*, low values indicate that the community is dominated by more resource-acquisitive species such as *Pinus*. Plant standing biomass and plant functional composition were negatively correlated (r=-0.57).

### Invertebrate herbivore and predator biomass

To characterize the invertebrate communities on the islands, we sampled invertebrates on each of the six dominant plant species on each island over a 10-day period in July 2016 (which is when plant productivity is at its peak). For each tree species (*Pinus, Betula* and *Picea*) on each island, we randomly selected five trees, and sampled one branch per tree. We collected insects from each branch using a beating sheet made of white fabric supported by a 1m^2^ frame, which was placed below the branch (Fig. S2). The chosen branches on the five trees were all of a similar size, and more or less covered the beating sheet (see SM1, Fig. S1). The branch was beaten with a stick (2.5cm diameter, 80cm long) eight times, and all dislodged arthropods were quickly collected off the sheet with an aspirator (Wade *et al*. 2006). The samples were pooled within each tree species on each island.

For each of the three shrub species (*V. myrtillus, V. vitis-idaea* and *Empetrum*) on each island, we sampled invertebrates with a leaf blower set to suction mode (Stihl SH86), on two randomly selected shrub patches of 50cm x 50cm each. For each shrub species we selected patches with at least 80% cover by the target species. The sampling area was covered with a frame 45cm high to prevent arthropods escaping and was immediately vacuumed for 100s at full power. Arthropods were intercepted in a gauze bag within the nozzle of the suction sampler and were then transferred to jars filled with ethanol (Fig. S3). Samples were pooled within each shrub species for each island. We recognize that one-time surveys might not fully characterise the invertebrate community, but as the vegetation period in northern Sweden is short (< 3 months) they should provide a reasonable estimate and enable unbiased comparisons among islands.

In the lab, invertebrates from each sample (trees and dwarf shrubs) were sorted to order or suborder level. Individuals of the sub-orders Auchenorrhyncha and Heteroptera, and the orders Coleoptera and Diptera, were identified to species level by expert entomologists, as some of these groups can contain both herbivorous and predatory species. We classified invertebrates to the trophic levels ‘herbivores’ and ‘predators’, and further divided herbivores into those with chewing versus sucking mouthparts (Table S2). Although ants (Formicidae) can also be predators, we did not consider them as predators in our data set as they get most of their food from aphid honeydew in this system (Rosengren & Sundström 1987; Domisch *et al*. 2009).

To obtain an estimate of herbivore and predator biomass on each tree and dwarf shrub species on each island, we measured the body length of all individuals using a digital microscope (LEICA DVM6), and used length-mass regressions from Sohlström et al., (2018) to estimate individual fresh body mass (Table S3). For each plant species on each island, we separately summed up the biomass of all individuals classified as predators, chewing herbivores and sucking herbivores.

To obtain a measure of total herbivore and predator biomass per island, for each plant species we calculated the invertebrate biomass per g aboveground standing plant biomass (dry mass). We used existing allometric equations for both trees (based on diameter at breast height and tree species, Marklund et al. 1988) and shrubs (based on vegetative ground cover, Wardle et al., 2003) to convert our sampling unit (five branches for trees, 0.5m^2^ ground cover for shrubs) to plant biomass (for details see SM2). We then multiplied the invertebrate biomass per g plant biomass by estimates of the standing biomass (g/m^2^, quantified in Wardle et al., 2003, 2012) for each of the tree and shrub species on that island. Finally, we summed up the values across all plant species per island, to get one estimate of invertebrate biomass for each island. We estimated total herbivore biomass, predator biomass, chewing herbivore biomass, and sucking herbivore biomass per m^2^ for each island in this way.

### Herbivore damage and impact on planted phytometer tree seedlings

We assessed herbivore damage and herbivore impact on the growth of planted seedlings of each of three tree species which we used as phytometers. We planted phytometers on each island and grew them with and without an insecticide treatment. The use of phytometers has the advantage that the genetic material is relatively homogenous across the study, and any variation in phytometer damage and performance reflects variation in the local herbivore community (Gibson 2015). We used broad-leaved species that occur in the region, i.e., *Betula pubescens, Salix caprea*, and *Alnus incana*. We chose broad-leaved trees in preference to conifers because it is much easier to measure leaf damage on them, and in preference to Ericaceous shrubs because it is very difficult to raise the latter from seeds or cuttings. We collected seeds from one mother tree of each of the three tree species on the mainland (Umeå, Sweden, Latitude: 63.808674, Longitude: 20.347868) to minimize genetic variation and potential pre-adaptation to island conditions. In February 2017, seeds of each species were germinated seedlings grown in pots. In June 2017, we planted 10 seedlings per tree species on each island. Seedlings were arranged in five blocks, each containing two similarly-sized individuals of each of the species. Half of the individuals were treated with a systemic insecticide (Calpyso, Bayer, active ingredient: Thiacloprid) three times during the growing season (June, July, August) in 2017 and 2018; the other half were sprayed with water. This insecticide has been used in previous herbivore exclusion experiments (e.g. Burkhart & Nentwig, 2008) and has been shown to not have non-target effects on plant growth (Lommen *et al*. 2018). We sprayed insecticide only on the individual tree seedlings and not on the surrounding vegetation.

In August 2018 we visually assessed mean herbivore damage on each seedling by inspecting all leaves per seedling and estimating the percentage leaf area removed by chewing herbivores. To get one value of herbivore damage per island, we only considered tree seedlings that had not been sprayed with insecticide. We averaged the percentage herbivore damage of those unsprayed seedlings per species and island, and then calculated the mean percentage herbivore damage across all three species.

We then assessed the impact of herbivores on plant biomass. We harvested all phytometer seedlings and assessed their aboveground dry weight. We calculated a log-response ratio comparing the biomass of tree seedlings with insecticide (reduced herbivory) to those without insecticide (ambient herbivory) as LRR_(herbivore impact)_ = log(Biomass with insecticide)/log(Biomass without insecticide). A LRR_(herbivore impact)_ of 0 would indicate that herbivores have no impact on seedling biomass, a LRR_(herbivore impact)_ >0 would indicate that herbivores have a negative impact. To get one value of herbivore impact per island, we averaged the LRR_(herbivore impact)_ per species on that island, and calculated the mean LRR_(herbivore impact)_ over all three species.

### Quantifying predation rate using plasticine caterpillars

The use of artificial models of prey has proved suitable for the standardized assessment of relative predation pressures by different predators such as birds, mammals and arthropods (Richards & Coley 2007; Fáveri *et al*. 2008). Artificial caterpillars have been shown to receive similar levels of attack as real caterpillars, providing a comparable measure of predation intensity among habitats (Richards & Coley 2007; Howe *et al*. 2009). On each island we set up a curved transect of 70m and marked it every 5m with a coloured bamboo stick (15 bamboo sticks totally) during July 2017. One meter to the left and right of each stick we attached a green plasticine caterpillar to a shrub using wire, resulting in 30 caterpillars per island. Caterpillars were each approximately 4mm x 30mm and were moulded from odourless, non-toxic, green coloured plasticine (Lewis Newplast) to imitate geometrid larvae, a common herbivore in this system. After two weeks we recorded whether caterpillars had been attacked by arthropods (small slits and scrape marks on the plasticine from mandibles), birds (characteristic beak marks) or small rodents (incisor marks, Low et al., 2014). We only used predation events by birds in our analysis, as we wanted to assess predation of a higher trophic group that was independent of our sampled invertebrate biomass. Rodent marks were very rare and were excluded. We used the number of caterpillars that were attacked by birds on each island as our response variable.

### Statistical analysis

We used structural equation models (SEMs) to identify the direct and indirect drivers of herbivore and predator biomass, bird predation and herbivore damage and impact. Our main question was whether invertebrate biomass depended on island size, and whether this effect was direct (path 3 in Fig. 1 and Table 1) or indirectly mediated by plant standing biomass (path 4) or plant functional composition (path 5). Additionally, we tested whether effects of island size and plants on herbivore biomass cascade up to predator biomass (path 6), and affect, or are affected by, bird predation (path 7). We tested models in which herbivore biomass is affected by predator biomass and bird predation (top-down influence) and in which predator biomass and bird predation are affected by herbivore biomass (bottom-up influence). These models were equivalent (same AIC values), but as herbivore biomass was positively related to bird predation and predator biomass, we consider the bottom-up model to be more ecologically plausible. We included an interaction term between predator biomass×standing plant biomass, to test whether the relationship between plant and herbivore biomass is stronger when predator biomass is low, and weaker when predator biomass is high (path 11 in Fig. 1 and Table 1), as predicted by the EEH hypothesis. Thus, our model contained a feedback loop, where herbivore biomass affects predator biomass and at the same time predator biomass affects herbivore biomass in interaction with plant biomass. Finally, we investigated the drivers of herbivore damage and herbivore impact on plant biomass (paths 13-16). We tested whether damage and impact are driven by herbivore biomass, but we also included a direct path from plant standing biomass and plant functional composition, as these variables might affect the functional composition of the herbivore community (which we did not measure) and might alter damage and impact independent of their effects on herbivore biomass. We included a direct path from predator biomass and bird predation rate to damage and impact for the same reason.

**Fig 1:**
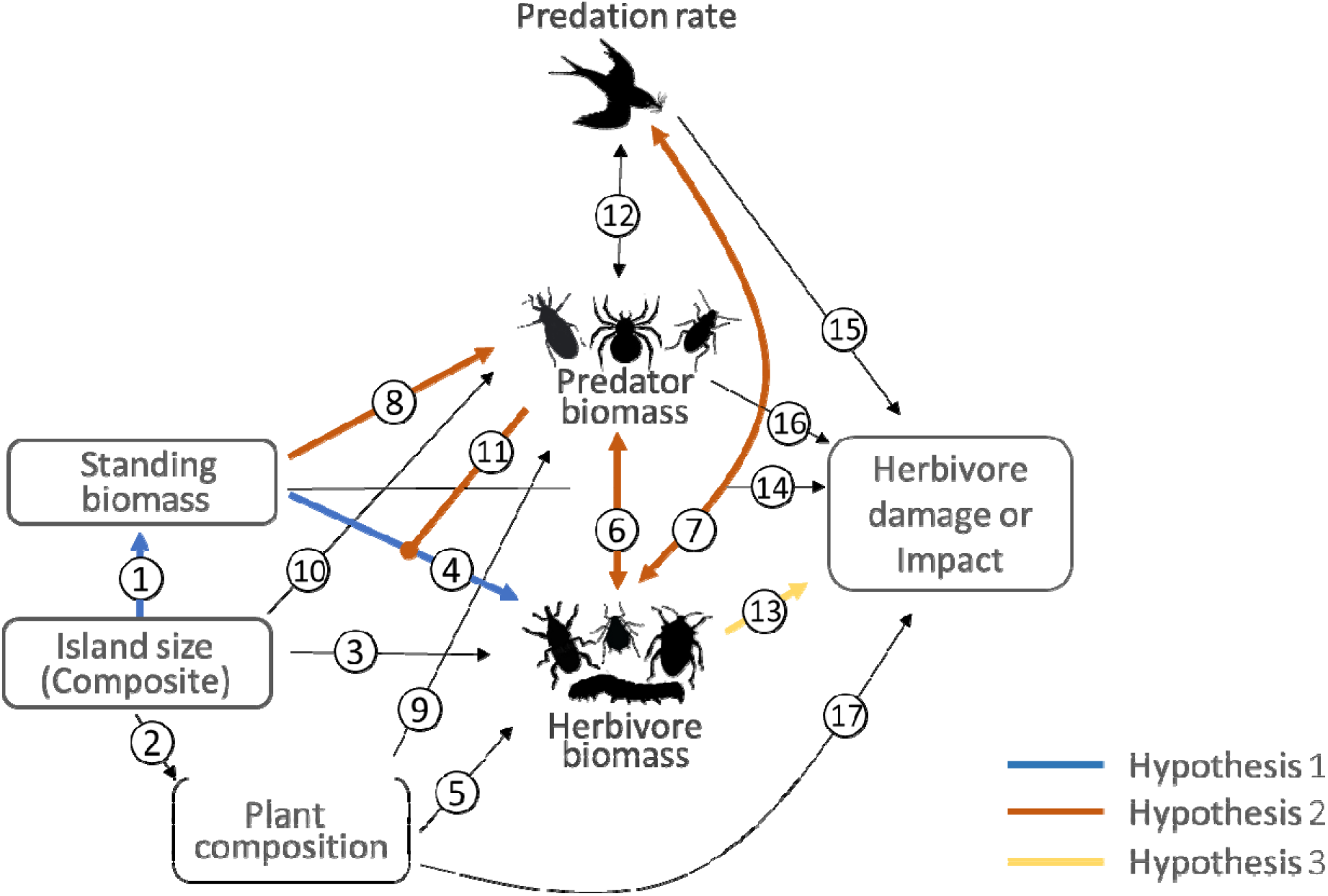
Schematic figure of the full SEM that was used to test the drivers of herbivore and predator biomass and herbivore damage. The path with the dot at one end (path 11) indicates a potential interactive effect of standing biomass and predator biomass on herbivore biomass. We tested all paths. The numbered paths and the hypotheses behind are explained in Table 1. Coloured paths in red, blue and yellow, respectively, correspond to the main hypotheses 1-3 that were tested.

We log-transformed island area, and the biomass of herbivores and predators. All variables were then standardized to comparable scales (Grace 2006). As we wanted to account for non-linear relationships between island size and the other variables, we included island size as a composite variable including the linear and the squared term of log-transformed island size. We ran different SEMs: one in which we included total herbivore biomass, one with only chewing herbivore biomass and one with only sucking herbivore biomass, to test whether the drivers of herbivore biomass differ between the different feeding guilds. To test whether the drivers of herbivore damage and impact are the same, we ran the model with total herbivore biomass, once with herbivore damage and once with impact. We removed non-significant paths from the model and tested whether their exclusion affected the overall model fit (i.e., whether it increased AIC). All analyses were performed using R (R Core Team 2017), and with the LAVAAN and lme4 package (Rosseel 2012; Bates *et al*. 2015).

## RESULTS

We collected 18979 invertebrate individuals, with a total biomass of 57450mg fresh weight. Of those, 3232 individuals (13009mg fresh weight) were herbivores (chewing herbivores: 546 individuals and 9737mg; sucking herbivores: 2686 individuals and 3272mg), and 10537 individuals (31261 mg) were predators (Table S2, Table S9).

In our SEM, island size strongly affected plant standing biomass, with larger islands having higher plant standing biomass (Fig. 2, Fig. 3A, Table S5). Island size also affected plant composition and was negatively related to PC1 (Fig. 2, Fig. 3B), meaning that larger islands were dominated by the most resource acquisitive species, *Pinus* and *V. myrtillus* and smaller islands by the most resource conservative species, *Picea* (Fig. S1).

**Fig. 2:**
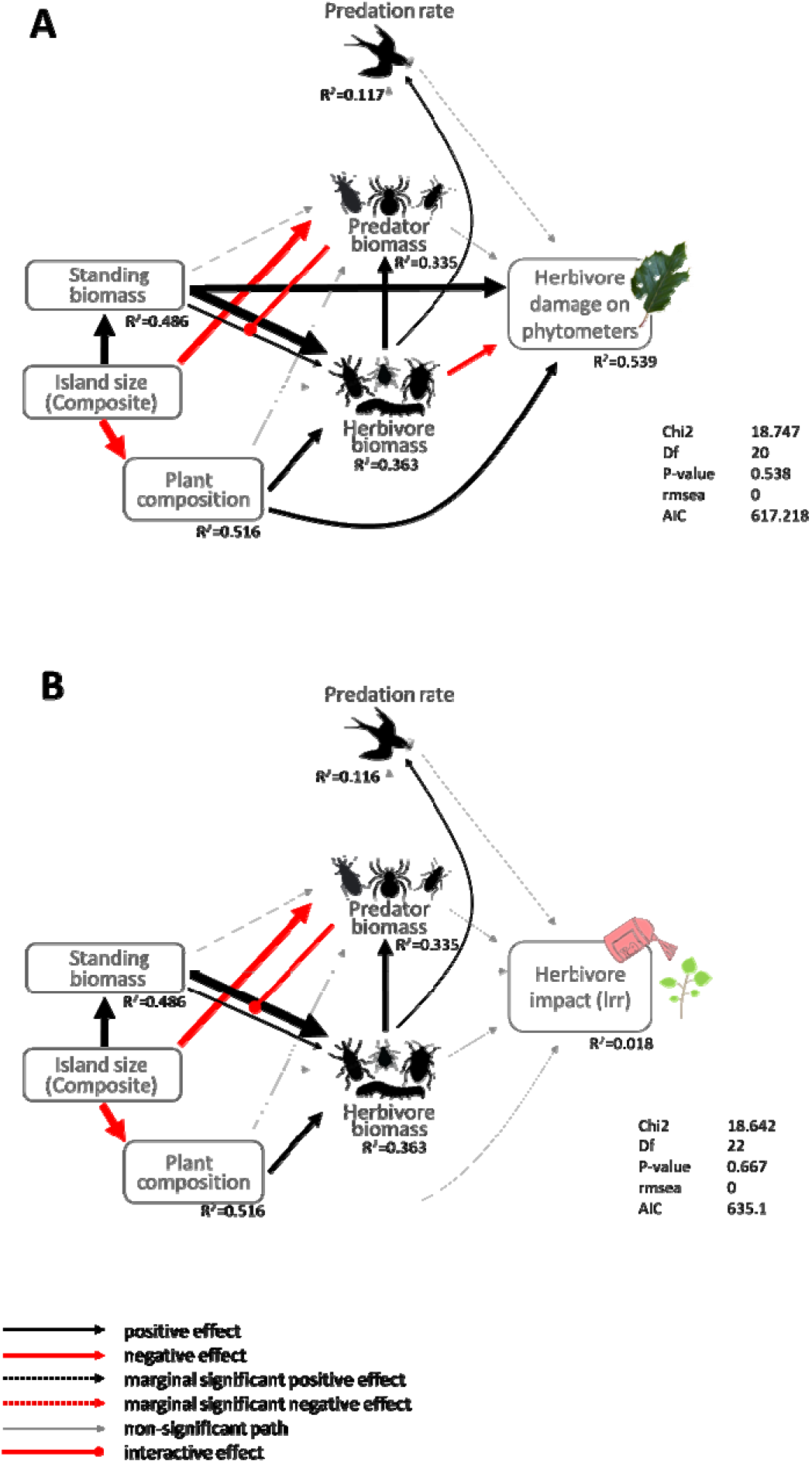
Results of structural equation models on the drivers and consequences of herbivore and predator biomass. A) investigates the drivers of herbivore damage on the phytometer tree seedlings, B) investigates the drivers of herbivore impact, which is the biomass of insecticide treated phytometer trees relative to the biomass of untreated phytometer trees (i.e., LRR_(herbivore impact)_). Black lines: significant positive effects at *P* = 0.05. Red lines: significant negative effects at *P* = 0.05. Grey lines: non-significant paths. The red line with the dot at the end indicates the interactive effect of standing biomass and predator biomass on herbivore biomass. Arrow width: relative strength of the path, see Table S5, S6.

**Fig. 3:**
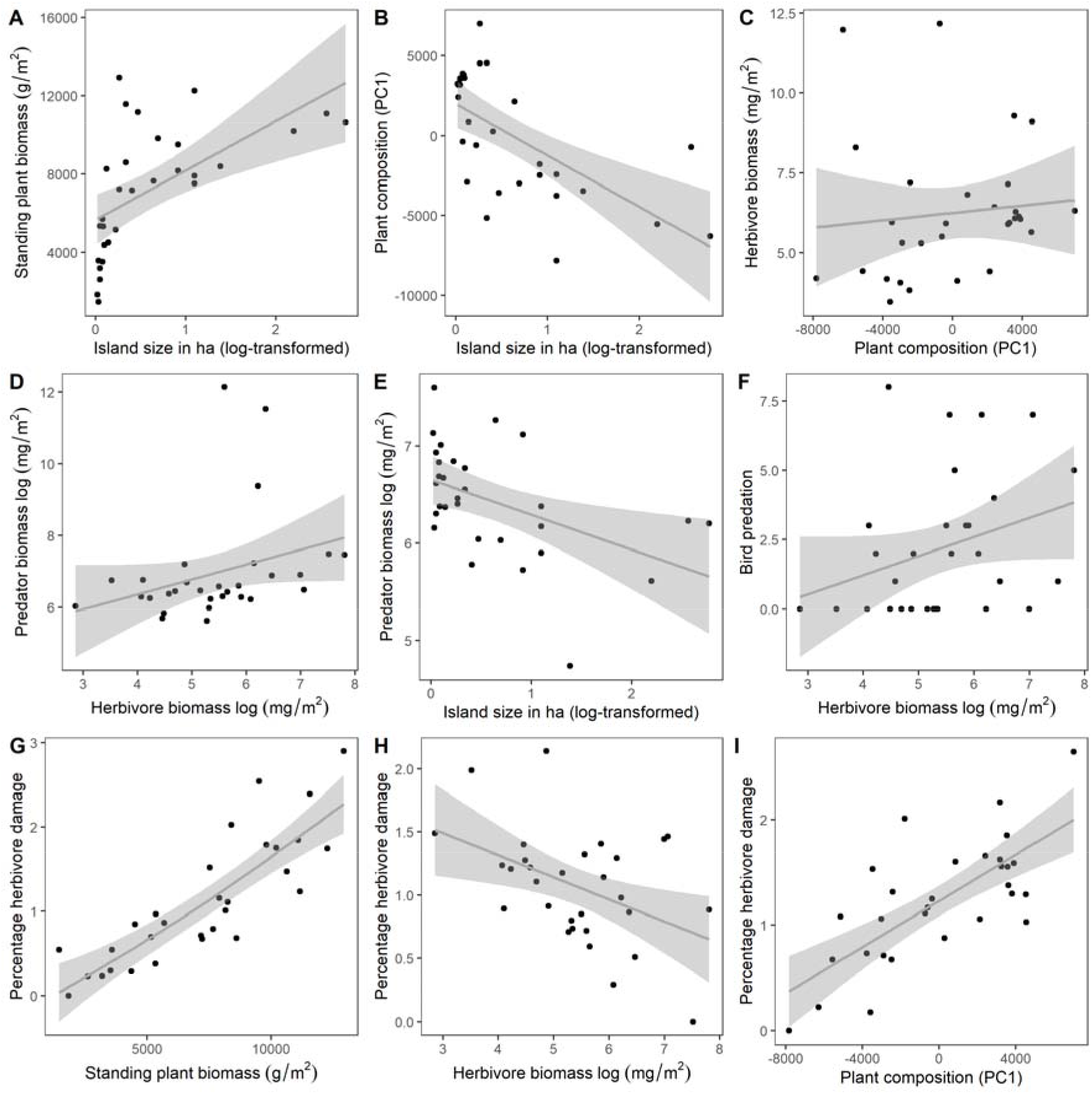
Partial plots of the SEM in Fig. 2A. Impact of significant predictor variables on standing plant biomass (A), plant composition (B), herbivore biomass (log-transformed) (C), predator biomass (log-transformed) (D,E), bird predation (F) and percentage herbivore damage on the phytometer seedlings (G-I), after removing all effects of all the other variables which are not plotted. Shaded areas represent 95% CI.

Herbivore biomass was strongly affected by bottom-up forces and increased with plant standing biomass (Fig. 2A). The strength of this relationship depended on predator biomass – when predator biomass was high, the relationship between herbivore biomass and plant standing biomass was weaker (as shown by the interactive effect of predator biomass and plant standing biomass on herbivore biomass in the SEM, Fig. 2, Fig. 3D). This agrees with the trophic flip predicted by the EEH. Herbivore biomass was also affected by plant species composition, and was higher in plant communities dominated by more resource conservative species (Fig. 2, Fig. 3C).

Predator biomass was higher on islands with a higher herbivore biomass (Fig. 2, Fig. 3D). This effect was mainly driven by chewing herbivores (Fig. 4, Table S7). Predator biomass was also higher on smaller islands (Fig. 2, Fig. 3F, Table S9). The predation rate by birds, as shown by the attack rate on plasticine caterpillars, was higher on islands with a high herbivore biomass (Fig. 2, Fig. 3G). This positive effect of herbivores on predation rate by birds was mainly driven by chewing herbivore biomass (Fig. 4, Table S7, S8).

**Fig. 4:**
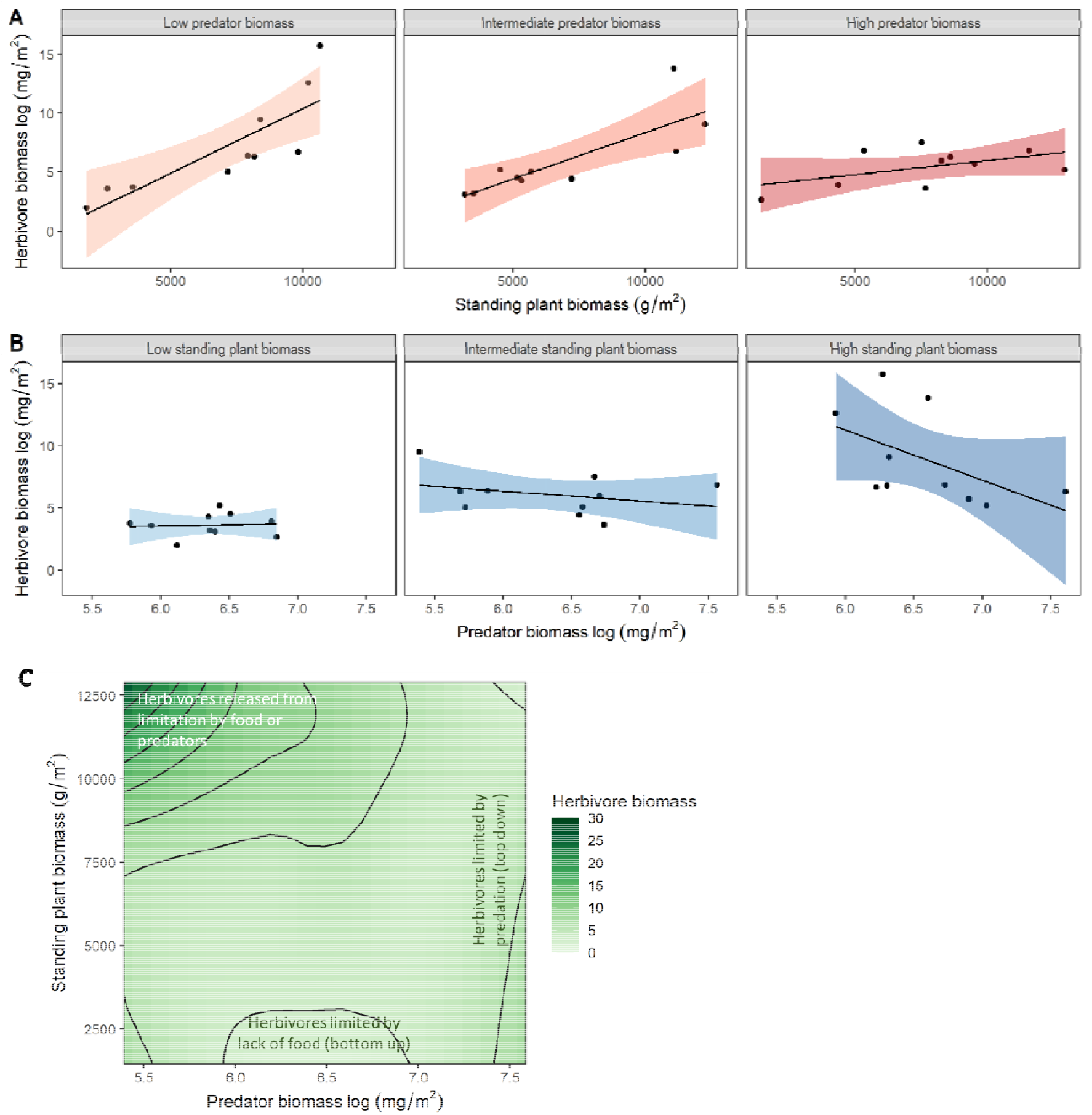
Partial plots of the SEM in Fig. 2A, showing the interactive effect of standing plant biomass and predator biomass on herbivore biomass, after removing all effects of all the other variables which are not plotted. A) Herbivore biomass increases strongly with plant standing biomass for low predator biomass, but less strongly for high predator biomass. B) Herbivore biomass is unresponsive to predator biomass for low standing plant biomass, but decreases for high plant biomass. For easier graphical visualisation, predator biomass (A) and plant standing biomass (B) has been binned into equal sized groups (low, medium and high), each group containing 10 islands (predator biomass: low [5.39 - 6.3], medium [6.3 - 6.63], high [6.63 - 7.61]; standing biomass: low [1460 - 5340], medium [5340 - 8460], high [8460 - 12900]). Shaded areas represent 95% CI. C) Shows the interactive effect of standing plant biomass and predator biomass on herbivore biomass in a contour plot, where dark colours indicate a high herbivore biomass.

**Fig. 5:**
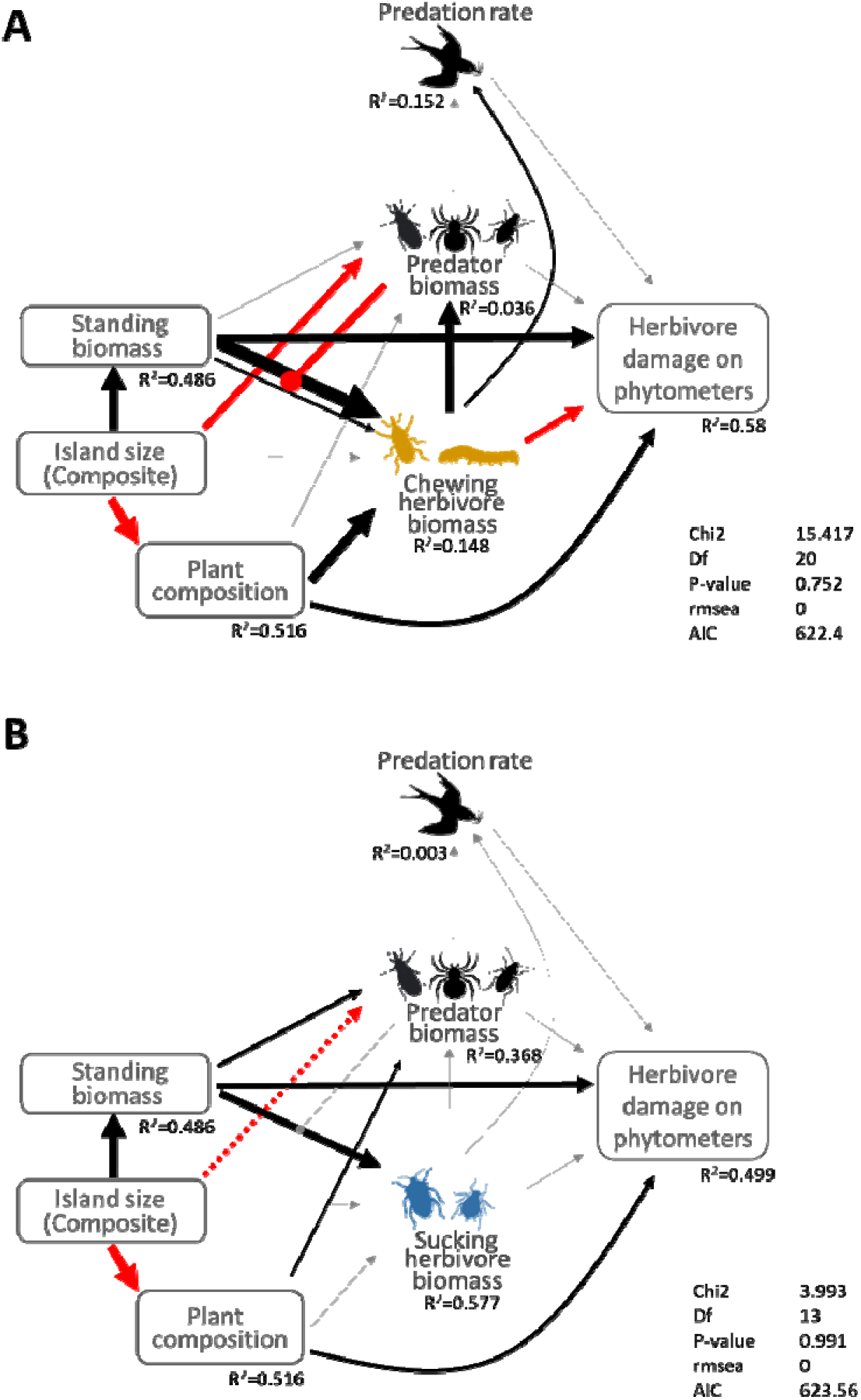
Structural equation models on the drivers and consequences of the biomass of A) chewing herbivores and B) sucking herbivores. Black lines: significant positive effects at *P* = 0.05. Red lines: significant negative effects at *P* = 0.05. The red line with the large dot at one end in panel A indicates the interactive effect of standing biomass and predator biomass on herbivore biomass. Grey lines: non-significant paths. The dashed red line in panel B indicates a marginally significant effect at P = 0.092. Thickness of line indicates strength of the path or correlation, see Table S7, S8.

The insecticide treatment was effective in reducing herbivore damage on the three phytometer species across all islands (SM3, Table S4). The damage on the unsprayed phytometer tree seedlings was highest on islands with a high standing plant biomass (Fig. 2A, Fig. 3H). This effect could not be explained by herbivore biomass (Fig. 2A). Plant composition also had strong direct effects on herbivore damage to the phytometer plants; phytometers on islands where resource conservative plant species dominated had the highest damage (Fig. 3J). In contrast, herbivore *impact* on phytometer biomass (comparing biomass of control and insecticide treated tree seedlings) was not affected by any of the tested drivers (Fig. 2B, Table S6).

## DISCUSSION

Our study across a retrogressive forested chronosequence shows that nutrient availability strongly controls invertebrate herbivore biomass when predators are rare, but that there is a switch from bottom-up to top-down control of herbivores when predators are common. This trophic flip is likely to contribute to the large variation we observe across herbivore impact studies (Coupe & Cahill 2003; Schädler *et al*. 2003; Jia *et al*. 2018), and can arise because invertebrate predators can respond to alternative energy channels independent of plant biomass. Our results further suggest that herbivore impact cannot necessarily be inferred from damage. We discuss these findings in the context of each of our three hypotheses to better understand the factors that regulates herbivore communities and their impacts on plants.

### Bottom-up versus top-down control of invertebrate herbivores

We found evidence that plant biomass controls invertebrate herbivore biomass: both plant and herbivore biomass declined with decreasing island size as nutrients become more limiting and plant biomass and productivity declines, which supports hypothesis 1. The cascading bottom-up effect of changes in plant biomass on higher trophic levels that we found is in line with empirical studies using anthropogenic fertility gradients and experiments in grasslands (e.g. Borer et al., 2012; Simons et al., 2014; Welti et al., 2020), as well as theory (Coley *et al*. 1985; McNaughton *et al*. 1989; Power 1992). However, few studies have explored invertebrate trophic responses to lengthy natural gradients of nutrient availability associated with ecosystem retrogression, and our results align with one previous study on total herbivore abundance in Hawaii (Gruner 2007), but not with another on a specialist weevil in our study system (Crutsinger *et al*. 2008). Specialist herbivores may respond differently to resource availability than would generalists or the entire herbivore community because they often use secondary compounds as cues to find their host plants or sequester chemicals for their own defense (Ali & Agrawal 2012), which could explain the discrepancy between our results and those of Crutsinger *et al*. (2008). Altogether, our results demonstrate that changes in plant biomass along natural soil fertility gradients involving retrogressive states cascade strongly up to affect higher trophic levels.

Despite the bottom-up effects of plant biomass on herbivore biomass, we also found evidence of top-down control of herbivores by predators. The strength of top-down control was not highest on the most productive islands as predicted by the EEH, but instead appeared to depend on predator biomass. This involved a switch from bottom-up control of herbivores when predators were few to top-down control when predators were common, in line with previous studies on vertebrates (Letnic & Ripple 2017) and two arthropod groups in grasslands (Welti *et al*. 2020). Thus, consistent with hypothesis 2, herbivore control flipped from bottom-up to top-down with increasing predator biomass. However, although plant standing biomass indirectly promoted predator biomass via the biomass of chewing herbivores, parts of hypothesis 2 was not confirmed as predators did not peak in abundance on islands with the highest plant biomass. Instead, predators were affected directly by island size, and their biomass was highest on the smallest (most nutrient limited) islands (Table S10). This may be because the predators in our system do not depend entirely on herbivores, and also use insects hatching from the surrounding lake water as prey (Jonsson *et al*. 2009). Small islands have a larger perimeter-to-area ratio, and thus on average a larger per-unit-area aquatic insect abundance than larger islands (Polis *et al*. 1997), supporting greater predator populations (Twining *et al*. 2016).

The partial independence of invertebrate predator biomass from plant and herbivore biomass that we found contrasts the strong relationship between plant and predator biomass found for vertebrate communities in several systems (e.g. Crête 1999; Letnic & Ripple 2017; Oksanen *et al*. 2020). However, it might apply more generally to invertebrate predators. For instance, many invertebrate predators also feed on pollinators which show contrasting responses to soil fertility compared to herbivores (Allan *et al*. 2014; Clough *et al*. 2014; Carvalheiro *et al*. 2020), or can benefit from cross-ecosystem flows of organisms such as between interconnected aquatic and terrestrial habitats (Gounand *et al*. 2018). In addition, insect predators might themselves be limited by predation from higher level predators (Gunnarsson & Wiklander 2015). Thus, the prediction of the EEH and hypothesis 2, that predator biomass is highest and herbivore impact reduced in productive systems, may not hold for invertebrates. However, the other prediction of the EEH, the trophic flip in herbivore control from bottom up when predators are few to top down when they are common, seems to be important in our system and partly confirms hypothesis 2. Our results suggest that switches between top-down and bottom-up control may indeed occur for invertebrate food webs but that these may be mediated more by changes in predator biomass than by cascading effects of changes in plant productivity.

### Herbivore damage is not equal to herbivore impact

Herbivore leaf damage of phytometers was highest on islands with the highest plant standing biomass, in support of the resource availability hypothesis, and hypothesis 3. Surprisingly however, phytometer leaf damage was not driven by total herbivore biomass or chewing herbivore biomass, but was directly promoted by standing plant biomass and dominance on the island of resource conservative plant species (see SEM in Fig. 2A, Hartley & Jones 1996). A direct effect of island plant biomass or composition on phytometer damage is unlikely, as increased plant biomass could not result in more damage without altering some aspect of the herbivore community. Instead, it is more plausible that variation in plant biomass and composition affected damage on the phytometers by altering the island’s herbivore community composition, which we did not measure. Moreover, damage was lower on phytometers growing on islands dominated by resource acquisitive species (*Pinus*). Large islands, with high plant biomass are dominated by resource acquisitive species, however, this result shows that after correcting for island size and plant biomass, phytometers on islands with a high abundance of *Pinus* actually suffered less damage. This may have occurred because our phytometers were all broadleaved species which might have suffered less attack by the herbivore community associated with *Pinus*.

Although herbivore leaf damage on the phytometers peaked on fertile islands, herbivore impact, i.e. the effect of herbivores on phytometer biomass, was unrelated to any predictor variables, or to leaf damage, in line with hypothesis 3. This brings into question the use of damage as an indicator of herbivore impact (Moles *et al*. 2011; Galmán *et al*. 2018). The lack of a relationship between damage and impact may have arisen from plants on more fertile islands being more tolerant of herbivore attack (Cronin *et al*. 2010). In addition, sap-sucking insects can strongly affect plant biomass, but their impact is hard to quantify visually. While some studies have suggested that the impact of herbivores on primary production varies as a function of productivity (Fraser & Grime 1997), the results of meta-analyses of insect exclusion studies in grasslands have been inconclusive (Coupe & Cahill 2003; Schädler *et al*. 2003; Jia *et al*. 2018). Thus, our results for tree seedling phytometers agree with findings from studies in grasslands that herbivore impact is not clearly linked to resource availability. However, exclusion experiments still remain rare, and more replicated exclusion experiments across contrasting ecosystems, like the one we performed here, or like those proposed by global research networks (e.g., the Bug-Network, bug-net.org) are key to further understanding the context-dependency of herbivore impacts on plant communities.

### Conclusion

We found a trophic flip in herbivore control, from bottom up, when predators were few, to top-down when predators were common, and which likely arose because predators were not affected by cascading effects of plant productivity but instead used additional energy channels. Our findings have several implications. Firstly, they highlight that classic theories of community regulation (e.g. Hairston *et al*. 1960; Oksanen *et al*. 1981), based on in situ productivity, can be incomplete when organisms forage across ecosystem boundaries. Such cross-ecosystem energy flow can lead to apparent competition between organisms from different systems, because increased resources in one system can increase predators, and hence top-down pressure on consumers, in the other system (Loreau *et al*. 2003). Secondly, our study calls for a better understanding of the drivers of herbivore impact. Herbivore community biomass may be a poor proxy of herbivore energy demand and impact (Ehnes *et al*. 2011; Barnes *et al*. 2018), and future studies that combine classical exclusion experiments with herbivore-impact assessments based on energy fluxes (Barnes *et al*. 2020) could help bring clarity to these relationships. Thirdly, our study highlights that retrogressive chronosequences can be useful model systems to study trophic relationships along *natural* soil fertility gradients. Given the large contribution of invertebrate herbivores to ecosystem functioning (Soliveres *et al*. 2016), and the ongoing changes in insect biomass currently occurring in terrestrial systems globally (van Klink *et al*. 2020), the need to improve our understanding of the drivers regulating insect biomass and impact is taking on a new importance.

## Acknowledgements

We thank Camille Martinez Almoyna and Rodrigo R. Granjel for help in the field and Elena Finkler for help with measuring invertebrates in the lab. The study was supported by funding for AK of the Swiss National Science foundation (P300PA_160991).

Supporting information see separate file

