## Supplementary material for "Plant-invertebrate interactions across a forested retrogressive chronosequence"

**Supporting information**

**Supplementary Methods:**

SM1: *Ensuring that the size of the branches sampled to collect invertebrates do not differ across the island size gradient*

To ensure that the branches we beat were of similar size, and did not vary in a biased manner across the island gradient, we estimated the length of each *Pinus* and *Picea* branch, and the dry biomass of leaves of each *Betula* branch, that we beat. To achieve this, we took photographs of each branch, together with the beating sheet. In the photographs of *Picea* and *Pinus* branches we also included a reference item of a known size (coloured in red, Fig. S2A, B).

For *Pinus* and *Picea* branches we then assessed the total branch length per photo, using image analysis (imageJ and photoshop). For *Betula* branches, we counted the number of leaves per photo. As the size of *Betula* leaves on smaller islands are smaller than on larger islands (Crutsinger et al., 2008), we additionally randomly sampled 30 leaves on each island and assessed their dry biomass, to obtain a value of mean dry biomass per leaf and island. We then estimated the dry biomass of *Betula* leaves per beaten branch by multiplying the number of leaves per branch (assessed with the photographs) by the mean leaf dry biomass of each island.

For each tree species we then obtained an average measure of branch length (*Pinus*, *Picea*) or an average measure of total leaf biomass per branch (*Betula*) for each island. We ran a linear model for each tree species to test whether the mean branch length (*Pinus*, *Picea*) or the leaf biomass per branch (*Betula*) that we beat differed among the three island size classes (small, medium, large).

There was no significant effect of island size at *P* = 0.05 on the estimated branch length for *Pinus* and *Picea* or on the dry biomass of *Betula* leaves per branch (*Pinus*: F_2,22_ = 0.575, P = 0.57; *Picea*: F_2,24_ = 0.63, P = 0.54; *Betula*: F_2,27_ = 2.66, P = 0.089). This means that there were no biases in our analyses that could be due to the size of branches that we beat varying with island size class for any of the three species.

SM2: *Converting invertebrate biomass caught per sampling unit (5 tree branches, 0.5m^2^ shrub cover) to a measure of invertebrate biomass per m^2^ for each island*

To convert our sampling unit (5 tree branches per tree species and island; 0.5 m^2^ of shrub cover) to g plant biomass we used published allometric equations. For the trees we used the allometric equations of Marklund et al. (1988), which presents separate equations for each tree species, to estimate branch biomass (dry weight basis) based on branch diameter. We assumed a diameter of all branches of 1.5 cm, and this resulted in calculated values per branch of 1300g dry mass for *Picea*, 630g for *Pinus*, and 410g for *Betula*. Thus, the total invertebrates caught on each island for each tree species were calculated as having occurred on 6500g dry weight for *Picea* (5 x 1300g), 3150g for *Pinus* and 2050g for *Betula*, allowing us to calculate the invertebrate biomass caught per g tree biomass.

For the dwarf shrubs, we used published allometric equations which convert aboveground vegetative cover for each of the three species to aboveground standing plant biomass (Wardle et al., 2003). We assumed that the sampled area of each shrub species had a cover of 95%, allowing us to calculate the aboveground standing biomass per shrub species per island on which invertebrates were sampled (141g for *V. vitis-ideae*, 137g for *V. myrtillus*, 184g for *Empetrum*). Again, this allowed us to calculate the invertebrate biomass caught per g shrub biomass.

Each of those values was then multiplied by estimates of the standing biomass (g dry mass per m^2^) of each of the six plant species per island, using standing biomass data for each island from Wardle et al. (2003, 2012). This resulted in a mean estimate of invertebrate biomass per square meter on an island, for each plant species. Finally, for each island we summed the values for all six plant species to obtain an estimate of invertebrate biomass (g) per m^2^ on each island.

SM3: *Effectiveness of the insecticide treatment in the phytometer approach*

We tested whether the insecticide treatment for the phytometer tree seedlings reduced the percentage herbivore damage. We used generalized linear mixed effect models, with treatment (with insecticide, without insecticide), phytometer species (*Alnus*, *Betula*, *Salix*), island size (small, medium, large) and their interactions as fixed effects, and island (30 levels) and block (5 levels) as random terms. We derived significance using likelihood-ratio tests comparing models with and without the factor of interest. We did all analyses in R version 4.0.3 (R Core Team, 2017).

The insecticide treatment significantly reduced herbivore damage on the three phytometer species across all islands (Table S4; estimate ± SE with insecticide = 0.34 ± 0.11, estimate ± SE without insecticide = 0.43±0.11). However, the reduction in herbivore damage did not result in a decrease in impact (estimate for LRR from an intercept only model with species, island and block as random terms = 0.0058, 95% CI [-0.049, 0.051]).

**Supplementary Tables**

Table S1: Measurements of selected ecosystem properties (mean values ± standard errors) measured at the whole island scale across the island size gradient. Data from Wardle and Zackrisson (2005) and Wardle et al. (1997, 2003, 2004, 2012).

| Ecosystem property |  | Island size |  |
| --- | --- | --- | --- |
|  | Small  (<0.1 ha) | Medium  (0.1 to 1.0 ha) | Large  (>1.0 ha) |
| Time since last fire (years) | 3250 ± 439 | 2180 ± 385 | 585 ± 233 |
| Island size (ha) | 0.058 ± 0.008 | 0.393 ± 0.073 | 4.8 ± 1.595 |
| Net primary productivity (g/m^2^/yr) | 159 ± 18 | 247 ± 12 | 256 ± 14 |
| Standing plant biomass (g/m^2^) | 3470 ± 470 | 8340 ± 877 | 9349 ± 485 |
| Humus C to N ratio | 32.9 ± 0.79 | 36.0 ± 1.17 | 40.4 ± 1.18 |
| Humus C to P ratio | 759 ± 30 | 687 ± 36 | 623 ± 20 |
| Mineral N (MIN) (µgN/g) | 25.3 ± 8.0 | 58.1 ± 9.2 | 38.2 ± 14.4 |
| Mineral P (µgP/g) | 24.4 ± 2.3 | 37.7 ± 4.3 | 43.6 ± 4.9 |

Table S2: List of Taxa. Individuals were identified to the taxonomic level necessary to determine their trophic level (i.e., herbivore or predator).

| **Taxa** | **Trophic group** |
| --- | --- |
| **Araneae** (all spiders) | Predator |
| **Blattodea** | Detritivore |
| **Ephemeroptera** | Aquatic (not considered) |
| **Trichoptera** | Aquatic (not considered) |
| **Megaloptera** | Aquatic (not considered) |
| **Plecoptera** | Aquatic (not considered) |
| **Gastropoda** (snails) | Herbivore |
| **Hymenoptera** - Formicidae (ants) | Omnivores |
| Hymenoptera - Parasitic wasps | Predator |
| Hymenoptera - Symphyta (Sawfly larvae) | Herbivore |
| **Lepidoptera** larvae | Herbivore |
| Lepidoptera - Geometridae larvae | Herbivore |
| **Neuroptera** - Chrysopidae | Predator |
| Neuroptera - Hemerobiidae | Predator |
| **Psocoptera** | Detritivore, Fungivore |
| **Coleoptera** - Cantharidae - Absidia schoenherri | Predator |
| Coleoptera - Cantharidae - Absidia schoenherri | Predator |
| Coleoptera - Cantharidae - Malthodes brevicollis | Predator |
| Coleoptera - Cantharidae - Malthodes brevicornis | Predator |
| Coleoptera - Cantharidae - Malthodes fuscus | Predator |
| Coleoptera - Cantharidae - Malthodes guttifer | Predator |
| Coleoptera - Cantharidae - Malthodes marginatus | Predator |
| Coleoptera - Cantharidae - Malthodes mysticus | Predator |
| Coleoptera - Cantharidae - Malthodes pumilus | Predator, Herbivore |
| Coleoptera - Cantharidae - Malthodes sp. | Predator, Herbivore |
| Coleoptera - Cantharidae - Malthods maurus | Predator |
| Coleoptera - Cantharidae - Podabrus alpinus | Predator |
| Coleoptera - Cantharidae - Podistra rufotestacea | Predator |
| Coleoptera - Cantharidae - Rhagonycha atra | Predator |
| Coleoptera - Carabidae - Dromius agilis | Predator |
| Coleoptera - Coccinellidae - Calvia quartordecimpunctata | Predator |
| Coleoptera - Coccinellidae - Coccinella hieroglyphica | Predator |
| Coleoptera - Coccinellidae - Scymnus redtenbacheri | Predator |
| Coleoptera - Coccinellidae - Scymnus sp. | Predator |
| Coleoptera - Cryptophagidae - Antherophagus alpinus | Fungivore |
| Coleoptera - Cryptophagidae - Cryptophagus subdepressus | Fungivore |
| Coleoptera - Curculionidae - Anoplus plantaris | Herbivore |
| Coleoptera - Curculionidae - Hylobius excavatus | Herbivore |
| Coleoptera - Curculionidae - Hylobius pinastri | Herbivore |
| Coleoptera - Curculionidae - Polydrusus fulvicornis | Herbivore |
| Coleoptera - Elateridae - Ampedus nigrinus | Detritivore |
| Coleoptera - Elateridae - Selatosomus impressus | Predator, Herbivore |
| Coleoptera - Scirtidae - Cyphon padi | Predator, Herbivore |
| Coleoptera - Scraptiidae - Anaspis arctica | Unknown |
| Coleoptera - Staphylinidae - Anthophagus omalinus | Predator |
| Coleoptera - Staphylinidae - Atheta sp. | Predator |
| Coleoptera - Staphylinidae - Mycetoporus splendidus | Predator |
| Coleoptera - Staphylinidae - Stenus clavicornis | Predator |
| Coleoptera - Staphylinidae - Stenus geniculatus | Predator |
| Coleoptera - Staphylinidae - Stenus ludyi | Predator |
| Coleoptera - Staphylinidae - Stenus palustris | Predator |
| **Diptera** - Agromyzidae | Herbivore |
| Diptera - Anthomyiidae | Herbivore |
| Diptera - Chamaemyiidae - Leucopis sensu lato sp. | Predator |
| Diptera - Chloropidae - Aphanotrigonum nigripes | Detritivore |
| Diptera - Chloropidae - Pseudopachychaeta ruficeps | Predator, Herbivore |
| Diptera - Cypselosomatidae - Pseudomyza atrimana | Detritivore |
| Diptera - Diastatidae - Diastata flavicosta | Detritivore, Fungivore |
| Diptera - Diastatidae - Diastata ornata | Detritivore, Fungivore |
| Diptera - Diastatidae - Diastata vagans | Detritivore, Fungivore |
| Diptera - Drosophilidae - Drosophila sp. | Predator, Herbivore |
| Diptera - Drosophilidae - Drosophila transversa | Predator, Herbivore |
| Diptera - Drosophilidae - Scaptomyza pallida | Herbivore |
| Diptera - Empiidae - Chelifera concinnicauda | Predator |
| Diptera - Empiidae - Chelifera sp. | Predator |
| Diptera - Empiidae - Rhamphomyia anomalina | Predator |
| Diptera - Empiidae - Rhamphomyia hybotina | Predator |
| Diptera - Fanniidae - Fannia postica | Detritivore, Fungivore |
| Diptera - Heleomyzidae - Suillia bicolor | Detritivore, Fungivore |
| Diptera - Hybotidae - Bicellaria austriaca | Predator |
| Diptera - Hybotidae - Bicellaria nigra | Predator |
| Diptera - Hybotidae - Euthyneura myrtilli | Predator |
| Diptera - Hybotidae - Platypalpus ecalceatus | Predator |
| Diptera - Hybotidae - Platypalpus nigritarsis | Predator |
| Diptera - Hybotidae - Platypalpus stigmatellus | Predator |
| Diptera - Hybotidae - Tachypeza nubila | Predator |
| Diptera - Lauxaniidae - Sapromyza hyalinata | Detritivore, Fungivore |
| Diptera - Muscidae | Predator, Detritivore |
| Diptera - Mycetophilidae | Detritivore, Fungivore |
| Diptera - Nematocera | Aquatic (not considered) |
| Diptera - Phoridae - Phoridae | Detritivore, Fungivore |
| Diptera - Platypezidae - Callomyia speciosa | Fungivore |
| Diptera - Scathophagidae - Scathophaga spurca | Herbivore |
| Diptera - Sciomyzidae - Ectinocera borealis | Predator |
| Diptera - Sphaeroceridae - Spelobia sp. | unknown |
| Diptera - Sphaeroceridae | unknown |
| **Hemiptera** - Heteroptera - Anthocoridae | Predator, Herbivore |
| Hemiptera - Heteroptera - Anthocoridae - Anthocoris nemorum | Predator, Herbivore |
| Hemiptera - Heteroptera - Anthocoridae - Anthocoris sp. | Predator, Herbivore |
| Hemiptera - Heteroptera - Anthocoridae - Tetraphleps bicuspis | Predator |
| Hemiptera - Sternorrhyncha - Aphididae | Herbivore |
| Hemiptera - Cicadomorpha - Cicadellidae - Colladonus torneellus | Herbivore |
| Hemiptera - Cicadomorpha - Cicadellidae - Idiodonus cruentatus | Herbivore |
| Hemiptera - Cicadomorpha - Cicadellidae - Macustus grisescens | Herbivore |
| Hemiptera - Cicadomorpha - Cicadellidae - Oncopis alni | Herbivore |
| Hemiptera - Cicadomorpha - Cicadellidae - Oncopis sp. | Herbivore |
| Hemiptera - Cicadomorpha - Cicadellidae - Oncopis subangulata | Herbivore |
| Hemiptera - Cicadomorpha - Cicadellidae - Oncopsis tristis | Herbivore |
| Hemiptera - Cicadomorpha - Cicadellidae - Wagneripteryx germari | Herbivore |
| Hemiptera - Heteroptera - Lygaeidae - Eremocoris abietis | Predator, Herbivore |
| Hemiptera - Heteroptera - Lygaeidae - Ligyrocoris sylvestris | Herbivore |
| Hemiptera - Heteroptera - Lygaeidae | Herbivore |
| Hemiptera - Heteroptera - Lygaeidae - Rhyparochromus pini | Herbivore |
| Hemiptera - Heteroptera - Lygaeidae - Scolopostethus sp. | Herbivore |
| Hemiptera - Heteroptera - Microphysidae - Loricula pselaphiformis | Predator |
| Hemiptera - Heteroptera - Miridae - Bothynotus pilosus | Herbivore |
| Hemiptera - Heteroptera - Miridae - Miridae (nymph) | Herbivore |
| Hemiptera - Heteroptera - Miridae - Oncotylus punctipes | Herbivore |
| Hemiptera - Heteroptera - Miridae - Plesiodema pinetella | Predator, Herbivore |
| Hemiptera - Heteroptera - Miridae - Psallus falleni | Herbivore |
| Hemiptera - Heteroptera - Miridae - Psallus salicis | Herbivore |
| Hemiptera - Auchenorrhyncha | Herbivore |
| Hemiptera - Cicadomorpha | Herbivore |
| Hemiptera - Heteroptera - Pentatomidae | Predator, Herbivore |
| Hemiptera - Sternorrhyncha - Psylloidea - Cacopsylla ambigua | Herbivore |
| Hemiptera - Sternorrhyncha - Psylloidea - Psylla betulae | Herbivore |
| Hemiptera - Sternorrhyncha - Psylloidea - Psylla betulaenanae | Herbivore |
| Hemiptera - Sternorrhyncha - Psylloidea - Psyllinae | Herbivore |
| Hemiptera - Heteroptera - Rhyparochromidae - Gastrodes abietum | Herbivore |
| Hemiptera - Heteroptera - Tingidae - Acalypta nigrina | Herbivore |
| Hemiptera - Heteroptera - Tingidae - Acalypta sp. | Herbivore |

Table S3: The length-mass regression parameter sources and model specifics for the invertebrates in this study. If not noted otherwise, regressions derived for temperate animals were preferred over those derived for tropical ones. All regressions resulted in fresh body masses.

| **Taxon** | **Regression reference** | **Regression name** |
| --- | --- | --- |
| Araneae | Sohlström et al. 2018 | LTR Araneae |
| Blattodea | Sohlström et al. 2018 | LTR Blattodea (tropical) |
| Coleoptera | Sohlström et al. 2018 | LTR Coleoptera |
| Coleoptera - Staphylinidae | Sohlström et al. 2018 | LTR Staphylinidae |
| Coleoptera - larvae | Sohlström et al. 2018 | LTR Coleoptera (larvae) |
| Diptera | Sohlström et al. 2018 | LTR Diptera |
| Ephemeroptera | Sohlström et al. 2018 | LR main |
| Trichoptera | Sohlström et al. 2018 | LR main |
| Plecoptera | Sohlström et al. 2018 | LR main |
| Megaloptera | Sohlström et al. 2018 | LR main |
| Gastropoda | Wardhaugh 2013;  Mercer et al. 2001 | L-dry mass and Mercer dry mass – fresh mass |
| Hemiptera | Sohlström et al. 2018 | LTR Hemiptera (other) |
| Hemiptera – Heteroptera | Sohlström et al. 2018 | LTR Hemiptera (Heteroptera) |
| Hemiptera juvenile | Sohlström et al. 2018 | LTR Hemiptera (juvenile |
| Hymenoptera | Sohlström et al. 2018 | LTR Hymenoptera 2 |
| Hymenoptera - Formicidae | Sohlström et al. 2018 | LTR Hymenoptera 1 |
| Lepidoptera | Sohlström et al. 2018 | LTR Lepidoptera |
| Lepidotpera larvae | Sohlström et al. 2018 | LTR Lepidoptera (larvae) |
| Neuroptera | Sohlström et al. 2018 | LTR Neuroptera |
| Psocodea | Sohlström et al. 2018 | LTR Psocoptera |

Table S4: Effects of phytometer species, insecticide treatment and island size class (small, medium, large) on percent herbivore foliar damage. We used linear mixed effect models and simplified models by removing non-significant terms from the model. We obtained significance by using likelihood-ratio tests comparing models with and without the factor of interest.

|  |  |  | ***Phytometer percent damage*** | | |
| --- | --- | --- | --- | --- | --- |
| ***Fixed terms*** |  |  | ***Df*** | ***Chi 2*** | ***P-value*** |
| Species |  |  | **2** | **56.122** | **<0.0001** |
| Treatment |  |  | **1** | **6.205** | **0.0127** |
| Island size class |  |  | 2 | 3.849 | 0.146 |
| Species x Treatment |  |  | 2 | 0.983 | 0.612 |
| Species x Island size class |  |  | 4 | 1.908 | 0.753 |
| Treatment x Island size class |  |  | 2 | 0.406 | 0.816 |
| Species x Treatment x Island size class |  |  | 4 | 8.922 | 0.063 |
| ***Random*** |  |  | *Variance* | *Stdev* |  |
| Island |  |  | 0.147 | 0.385 |  |
| Block |  |  | <0.0001 | <0.0001 |  |

Table S5: Paths, correlations and estimates for the drivers of total herbivore biomass, predator biomass and phytometer damage. The table corresponds to Fig. 2A. Standing plant biomass has been obtained from Wardle et al. 2003, 2012 and describes the total plant standing biomass of the six dominant plant species (i.e. *Pinus*, *Betula*, *Picea*, *V. myrtillus*, *V. vitis-idaea* and *Empetrum*) per square meter. Island size has been included as a composite variable of the linear and the squared term of the log-transformed island size (in ha) to also account for non-linear relationships between island size and the other variables.

| **Path** |  |  |  |  |  |  |
| --- | --- | --- | --- | --- | --- | --- |
| *Response* | *Predictor* |  | *Estimate* | *SE* | *z-value* | *p-value* |
| ***Composites*** |  |  |  |  |  |  |
| Island size (composite) | Island size |  | 1 |  |  |  |
|  | (Island size)^2^ |  | -0.303 | 0.04 | -7.626 | 0 |
| ***Regressions*** |  |  |  |  |  |  |
| Standing plant biomass | Island size (composite) | | **1.136** | **0.221** | **5.133** | **<0.0001** |
| Plant composition | Island size (composite) | | **-1.175** | **0.217** | **-5.422** | **<0.0001** |
| Herbivore biomass | Island size (composite) | | -0.691 | 0.579 | -1.194 | 0.233 |
|  | Plant composition | | **0.77** | **0.274** | **2.807** | **0.005** |
|  | Standing plant biomass | | **1.486** | **0.361** | **4.118** | **<0.0001** |
|  | Predator biomass | | -0.652 | 0.335 | -1.944 | 0.052 |
|  | Standing biomass x Predator biomass | | **-0.688** | **0.244** | **-2.82** | **0.005** |
| Predator biomass | Island size (composite) | | **-1.053** | **0.297** | **-3.542** | **<0.0001** |
|  | Standing plant biomass | | -0.204 | 0.582 | -0.351 | 0.726 |
|  | Plant composition | | 0.23 | 0.247 | 0.928 | 0.353 |
|  | Herbivore biomass | | **0.896** | **0.204** | **4.398** | **<0.0001** |
| Phytometer damage | Herbivore biomass | | **-0.341** | **0.167** | **-2.098** | **0.036** |
|  | Standing plant biomass | | **1.126** | **0.212** | **5.301** | **<0.0001** |
|  | Bird predation | | 0.147 | 0.132 | 1.11 | 0.267 |
|  | Predator biomass | | 0.072 | 0.15 | 0.476 | 0.634 |
|  | Plant composition | | **0.737** | **0.166** | **4.441** | **<0.0001** |
| Bird predation | Herbivore biomass | | **0.312** | **0.156** | **1.995** | **0.046** |
|  | Predator biomass | | -0.123 | 0.17 | -0.728 | 0.467 |
| ***Covariances*** |  |  |  |  |  |  |
| Island size | (Island size)^2^ |  | 1.53 | 0.437 | 3.502 | <0.0001 |
| Phytometer damage | Bird predation | | 0.128 | 0.12 | 1.06 | 0.286 |
| ***Variances:*** |  |  |  |  |  |  |
| Island size composite |  |  | 0 |  |  |  |
| Standing plant biomass |  |  | 0.497 | 0.128 | 3.873 | <0.0001 |
| Plant composition |  |  | 0.468 | 0.121 | 3.873 | <0.0001 |
| Herbivore biomass |  |  | 0.758 | 0.348 | 2.18 | 0.029 |
| Predator biomass |  |  | 0.823 | 0.253 | 3.248 | 0.001 |
| Phytometer damage |  |  | 0.475 | 0.123 | 3.873 | <0.0001 |
| Bird predation |  |  | 0.873 | 0.225 | 3.873 | <0.0001 |
| Island size |  |  | 0.967 | 0.25 | 3.873 | <0.0001 |
| (Island size)^2^ |  |  | 3.501 | 0.904 | 3.873 | <0.0001 |

Table S6: Paths, correlations and estimates for the drivers of total herbivore biomass, predator biomass and herbivore *impact* on phytometer seedlings. The table corresponds to Fig. 2B. Standing plant biomass has been obtained from Wardle et al. (2003, 2012) and describes the total plant standing biomass of the six dominant plant species (i.e. *Pinus*, *Betula*, *Picea*, *V. myrtillus*, *V. vitis-idaea* and *Empetrum*) per square meter. Island size has been included as a composite variable of the linear and the squared term of the log-transformed island size (in ha) to also account for non-linear relationships between island size and the other variables. Herbivore impact has been measured as the log response ratio (lrr) of the biomass of the phytometer seedlings with insecticide / the biomass of the seedlings without insecticide.

| **Path** |  |  |  |  |  |  |
| --- | --- | --- | --- | --- | --- | --- |
| *Response* | *Predictor* |  | *Estimate* | *SE* | *z-value* | *p-value* |
| ***Composites*** |  |  |  |  |  |  |
| Island size (composite) | Island size |  | 1 |  |  |  |
|  | (Island size)^2^ |  | -0.3 | 0.039 | -7.62 | <0.0001 |
| ***Regressions*** |  |  |  |  |  |  |
| Standing plant biomass | Island size (composite) |  | **1.136** | **0.221** | **5.133** | **<0.0001** |
| Plant composition | Island size (composite) |  | **-1.17** | **0.216** | **-5.43** | **<0.0001** |
| Herbivore biomass | Island size (composite) |  | -0.691 | 0.579 | -1.19 | 0.233 |
|  | Plant composition |  | **0.77** | **0.274** | **2.807** | **0.005** |
|  | Standing plant biomass |  | **1.486** | **0.361** | **4.118** | **<0.0001** |
|  | Predator biomass |  | -0.652 | 0.335 | -1.94 | 0.052 |
|  | Standing biomass x Predator biomass | | **-0.687** | **0.244** | **-2.82** | **0.005** |
| Predator biomass | Island size (composite) |  | **-1.053** | **0.297** | **-3.54** | **<0.0001** |
|  | Standing plant biomass |  | -0.204 | 0.582 | -0.35 | 0.726 |
|  | Plant composition |  | 0.23 | 0.247 | 0.928 | 0.353 |
|  | Herbivore biomass |  | **0.896** | **0.204** | **4.398** | **<0.0001** |
| Herbivore impact (lrr) | Herbivore biomass |  | -0.033 | 0.164 | -0.2 | 0.84 |
|  | Standing plant biomass |  | 0.144 | 0.302 | 0.477 | 0.63 |
|  | Bird predation |  | -0.028 | 0.19 | -0.15 | 0.881 |
|  | Predator biomass |  | -0.007 | 0.214 | -0.03 | 0.975 |
|  | Plant composition |  | 0.134 | 0.181 | 0.74 | 0.459 |
| Bird predation | Herbivore biomass |  | **0.31** | **0.156** | **1.987** | **0.047** |
|  | Predator biomass |  | -0.123 | 0.169 | -0.73 | 0.467 |
| ***Covariances*** |  |  |  |  |  |  |
| Island size | (Island size)^2^ |  | 1.53 | 0.437 | 3.502 | <0.0001 |
| Herbivore impact (lrr) | Bird predation |  | -0.053 | 0.166 | -0.32 | 0.751 |
| ***Variances:*** |  |  |  |  |  |  |
| Island size (composite) |  |  | 0 | 0 | 0 |  |
| Standing plant biomass |  |  | 0.497 | 0.128 | 3.873 | <0.0001 |
| Plant composition |  |  | 0.468 | 0.121 | 3.873 | <0.0001 |
| Herbivore biomass |  |  | 0.758 | 0.348 | 2.18 | 0.029 |
| Predator biomass |  |  | 0.823 | 0.253 | 3.248 | 0.001 |
| Herbivore impact (lrr) |  |  | 0.95 | 0.245 | 3.873 | <0.0001 |
| Bird predation |  |  | 0.873 | 0.225 | 3.873 | <0.0001 |
| Island size |  |  | 0.967 | 0.25 | 3.873 | <0.0001 |
| (Island size)^2^ |  |  | 3.501 | 0.904 | 3.873 | <0.0001 |

Table S7: Paths, correlations and estimates for the drivers of chewing herbivore biomass, predator biomass and phytometer damage. The table corresponds to Fig. 4A. Standing plant biomass has been obtained from Wardle et al. (2003, 2012) and describes the total plant standing biomass of the six dominant plant species (i.e. *Pinus*, *Betula*, *Picea*, *V. myrtillus*, *V. vitis-idaea* and *Empetrum*) per square meter. Island size has been included as a composite variable of the linear and the squared term of the log-transformed island size (in ha) to also account for non-linear relationships between island size and the other variables.

| **Path** |  |  |  |  |  |  |
| --- | --- | --- | --- | --- | --- | --- |
| *Response* | *Predictor* |  | *Estimate* | *SE* | *z-value* | *p-value* |
| ***Composites*** |  |  |  |  |  |  |
| Island size (composite) | Island size |  | 1 | 1.656 | 1.628 |  |
|  | (Island size)^2^ |  | -0.299 | 0.041 | -7.268 | 0 |
| ***Regressions*** |  |  |  |  |  |  |
| Standing plant biomass | Island size (composite) |  | **1.135** | **0.222** | **5.115** | **<0.0001** |
| Plant composition | Island size (composite) |  | **-1.169** | **0.216** | **-5.404** | **<0.0001** |
| Chewing herbivore biomass | Island size (composite) |  | -1 | 0.765 | -1.307 | 0.191 |
|  | Plant composition |  | **1.204** | **0.429** | **2.804** | **0.005** |
|  | Standing plant biomass |  | **1.692** | **0.513** | **3.296** | **0.001** |
|  | Predator biomass |  | **-1.163** | **0.519** | **-2.241** | **0.025** |
|  | Standing biomass x Predator biomass | | **-0.985** | **0.367** | **-2.683** | **0.007** |
| Predator biomass | Island size (composite) |  | **-0.894** | **0.349** | **-2.56** | **0.01** |
|  | Standing plant biomass |  | 0.135 | 0.459 | 0.294 | 0.769 |
|  | Plant composition |  | 0.059 | 0.336 | 0.176 | 0.861 |
|  | Chewing herbivore biomass |  | **1.047** | **0.277** | **3.776** | **<0.0001** |
| Phytometer damage | Chewing herbivore biomass |  | **-0.384** | **0.137** | **-2.805** | **0.005** |
|  | Standing plant biomass |  | **1.083** | **0.17** | **6.362** | **<0.0001** |
|  | Bird predation |  | 0.186 | 0.126 | 1.478 | 0.139 |
|  | Predator biomass |  | 0.033 | 0.135 | 0.246 | 0.806 |
|  | Plant composition |  | **0.782** | **0.154** | **5.092** | **<0.0001** |
| Bird predation | Chewing herbivore biomass |  | **0.357** | **0.154** | **2.323** | **0.02** |
|  | Predator biomass |  | -0.114 | 0.156 | -0.73 | 0.466 |
| ***Covariances*** |  |  |  |  |  |  |
| Island size | (Island size)^2^ |  | 1.53 | 0.437 | 3.502 | <0.0001 |
| Phytometer damage | Bird predation |  | 0.157 | 0.113 | 1.382 | 0.167 |
| ***Variances:*** |  |  |  |  |  |  |
| Island size (composite) |  |  | 0 |  |  |  |
| Standing plant biomass |  |  | 0.497 | 0.128 | 3.873 | <0.0001 |
| Plant composition |  |  | 0.468 | 0.121 | 3.873 | <0.0001 |
| Chewing herbivore biomass |  |  | 1.366 | 0.757 | 1.805 | 0.071 |
| Predator biomass |  |  | 1.257 | 0.489 | 2.571 | 0.01 |
| Phytometer damage |  |  | 0.429 | 0.111 | 3.873 | <0.0001 |
| Bird predation |  |  | 0.843 | 0.218 | 3.873 | <0.0001 |
| Island size |  |  | 0.967 | 0.25 | 3.873 | <0.0001 |
| (Island size)^2^ |  |  | 3.501 | 0.904 | 3.873 | <0.0001 |

Table S8: Paths, correlations and estimates for the drivers of sucking herbivore biomass, predator biomass and phytometer damage. The table corresponds to Fig. 4B. Standing plant biomass has been obtained from Wardle et al. (2003, 2012) and describes the total plant standing biomass of the six dominant plant species (i.e. *Pinus*, *Betula*, *Picea*, *V. myrtillus*, *V. vitis-idaea* and *Empetrum*) per square meter. Island size has been included as a composite variable of the linear and the squared term of the log-transformed island size (in ha) to also account for non-linear relationships between island size and the other variables.

| **Path** |  |  |  |  |  |  |
| --- | --- | --- | --- | --- | --- | --- |
| *Response* | *Predictor* |  | *Estimate* | *SE* | *z-value* | *p-value* |
| ***Composites*** |  |  |  |  |  |  |
| Island size (composite) | Island size |  | 1 | 1.65 | 1.622 |  |
|  | (Island size)^2^ |  | -0.296 | 0.042 | -7.035 | <0.0001 |
| ***Regressions*** |  |  |  |  |  |  |
| Standing plant biomass | Island size (composite) |  | **1.131** | **0.222** | **5.099** | **<0.0001** |
| Plant composition | Island size (composite) |  | **-1.165** | **0.216** | **-5.39** | **<0.0001** |
| Sucking herbivore biomass | Island size (composite) |  | -0.379 | 0.269 | -1.406 | 0.16 |
|  | Plant composition |  | -0.0001 | 0.178 | -0.002 | 0.998 |
|  | Standing plant biomass |  | **0.904** | **0.166** | **5.457** | **<0.0001** |
|  | Predator biomass |  | -0.141 | 0.135 | -0.1048 | 0.295 |
|  | Standing biomass x Predator biomass | | -0.066 | 0.148 | -0.442 | 0.659 |
| Predator biomass | Island size (composite) |  | -0.706 | 0.419 | -1.685 | 0.092 |
|  | Standing plant biomass |  | **0.627** | **0.207** | **3.033** | **0.002** |
|  | Plant composition |  | **0.428** | **0.213** | **2.008** | **0.045** |
|  | Sucking herbivore biomass |  | 0.122 | 0.699 | 0.174 | 0.862 |
| Phytometer damage | Sucking herbivore biomass |  | 0.125 | 0.199 | 0.628 | 0.53 |
|  | Standing plant biomass |  | **0.805** | **0.155** | **5.198** | **<0.0001** |
|  | Bird predation |  | 0.01 | 0.133 | 0.078 | 0.938 |
|  | Predator biomass |  | -0.002 | 0.158 | -0.015 | 0.988 |
|  | Plant composition |  | **0.63** | **0.155** | **4.073** | **<0.0001** |
| Bird predation | Sucking herbivore biomass |  | 0.052 | 0.182 | 0.286 | 0.775 |
|  | Predator biomass |  | -0.021 | 0.179 | -0.118 | 0.906 |
| ***Covariances*** |  |  |  |  |  |  |
| Island size | (Island size)^2^ |  | 1.53 | 0.437 | 3.502 | 0 |
| Mean damage | Bird predation |  | 0.014 | 0.129 | 0.105 | 0.916 |
| Predator biomass | Bird predation |  | 0.038 | 0.143 | 0.262 | 0.794 |
| Predator biomass | Phytometer damage |  | 0.014 | 0.105 | 0.134 | 0.894 |
| ***Variances:*** |  |  |  |  |  |  |
| Island size (composite) |  |  | 0 | 0 | 0 |  |
| Standing plant biomass |  |  | 0.497 | 0.128 | 3.873 | <0.0001 |
| Plant composition |  |  | 0.468 | 0.121 | 3.873 | <0.0001 |
| Chewing herbivore biomass |  |  | 0.409 | 0.106 | 3.873 | <0.0001 |
| Predator biomass |  |  | 0.639 | 0.165 | 3.873 | <0.0001 |
| Phytometer damage |  |  | 0.521 | 0.134 | 3.873 | <0.0001 |
| Bird predation |  |  | 0.964 | 0.249 | 3.873 | <0.0001 |
| Island size |  |  | 0.967 | 0.25 | 3.873 | <0.0001 |
| (Island size)^2^ |  |  | 3.501 | 0.904 | 3.873 | <0.0001 |

Table S9: Estimated (mean values ± standard errors) invertebrate biomass in mg fresh weight per m^2^ island (see SM2), and other measured variables for the three island size classes, and F_2,27_ and P-values from an ANOVA with island size class as the explanatory variable. Herbivore impact was assessed based on the log response ratio of the biomass of phytometer trees with insecticide / biomass without insecticide. Bird predation indicates the number of plasticine caterpillars that had been attacked by birds (30 caterpillars per island).

| Measured variable | Island size | | |  |  |
| --- | --- | --- | --- | --- | --- |
|  | Small | Medium | Large |  |  |
|  | (<0.1 ha) | (0.1 to 1.0 ha) | (>1.0 ha) |  |  |
|  | mean ± SE | mean ± SE | mean ± SE | F-Value | P-Value |
| Herbivore biomass | 332 ± 237 | 485 ± 176 | 442 ± 99 | 3.707 | **0.038** |
| Predator biomass | 739 ± 147 | 848 ± 147 | 535 ± 76 | 2.103 | 0.142 |
| Chewing herbivore biomass | 303 ± 235 | 337 ± 169 | 330 ± 110 | 0.721 | 0.496 |
| Sucking herbivore biomass | 28 ± 7 | 147 ± 43 | 111 ± 25 | 11.164 | **0.0003** |
| Phytometer damage [%] | 0.201 ± 0.044 | 0.538 ± 0.26 | 0.507 ± 0.168 | 1.06 | 0.359 |
| Herbivore impact (lrr) | 0.025 ± 0.051 | -0.032 ± 0.063 | 0.004 ± 0.022 | 0.354 | 0.705 |
| Bird predation | 1.7 ± 0.87 | 2.5 ± 0.703 | 2.4 ± 0.884 | 0.28 | 0.758 |

Table S9: The biomass of caught invertebrates (mean values ± standard errors) in mg fresh weight per plant species, and other measured variables for the three island size classes, and F_2,27_ and P-values from an ANOVA with island size class as the explanatory variable. Invertebrate biomass on trees are per 5 branches per species and islands, for shrubs per 0.5m^2^ per species and island. ANOVAs for invertebrate biomass were run separately for each plant species. Herbivore impact has been assessed based on the log response ratio of the biomass of phytometer trees with insecticide to biomass without insecticide. Bird predation indicates the number of plasticine caterpillars that had been attacked by birds (30 caterpillars per island).

| Measured variable | | Island size | | |  |  |
| --- | --- | --- | --- | --- | --- | --- |
|  |  | Small | Medium | Large |  |  |
|  |  | (<0.1 ha) | (0.1 to 1.0 ha) | (>1.0 ha) |  |  |
|  |  | mean ± SE | mean ± SE | mean ± SE | F-Value | P-Value |
| Herbivore biomass on | |  |  |  |  |  |
|  | *Betula* | 50.4±28.0 | 75.0±16.7 | 81.9±55.8 | 1.569 | 0.267 |
|  | *Pinus* | 37.6±19.0 | 28.1±8.22 | 81.6±28.1 | 2.337 | 0.12 |
|  | *Picea* | 149.0±91.2 | 77.7±35.3 | 40.5±15.6 | 0.9339 | 0.407 |
|  | *V. vitis-idaea* | 160.0±136.0 | 123.0±62.1 | 107.0±42.5 | 0.518 | 0.602 |
|  | *V. myrtillus* | 47.4±18.1 | 54.9±20.1 | 46.6±15.3 | 0.471 | 0.629 |
|  | *Empetrum* | 91.9±77.1 | 15.5±4.63 | 81.7±41.8 | 0.558 | 0.578 |
| Predator biomass on | |  |  |  |  |  |
|  | *Betula* | 184.0±40.1 | 85.6±31.5 | 100.0±21.9 | 2.177 | 0.133 |
|  | *Pinus* | 174.3±77.5 | 150.2±35.1 | 74.0±16.3 | 1.231 | 0.3113 |
|  | *Picea* | 607.4±149.1 | 322.1±82.6 | 223.0±32.1 | 3.014 | **0.068** |
|  | *V. vitis-idaea* | 294.0±58.8 | 169.1±28.5 | 109.5±24.8 | 8.208 | **0.002** |
|  | *V. myrtillus* | 229.0±41.4 | 202.8±27.4 | 86.6±14.5 | 7.304 | **0.003** |
|  | *Empetrum* | 134.0±66.9 | 117.0±22.8 | 75.9±10.5 | 2.41 | 0.109 |
| Chewing herbivore biomass on | |  |  |  |  |  |
|  | *Betula* | 35.7±27.1 | 13.7±9.35 | 46.2±38.8 | 0.291 | 0.75 |
|  | *Pinus* | 14.7±12.6 | 1.98±1.98 | 49.1±31 | 1.601 | 0.224 |
|  | *Picea* | 123.0±90.7 | 38.4±32.7 | 26.8±15.2 | 0.746 | 0.485 |
|  | *V. vitis-idaea* | 155.2±136.1 | 116.3±61.7 | 98.9±42.5 | 0.36 | 0.701 |
|  | *V. myrtillus* | 41.3±17.9 | 40.1±18.3 | 29.3±14.9 | 0.103 | 0.902 |
|  | *Empetrum* | 87.2±77.4 | 8.74±4.83 | 70.4±42.7 | 0.655 | 0.527 |
| Sucking herbivore biomass on | |  |  |  |  |  |
|  | *Betula* | 14.7±3.5 | 61.3±15.4 | 35.7±18.3 | 3.991 | **0.03** |
|  | *Pinus* | 22.9±8.24 | 26.1±6.99 | 32.4±8.93 | 0.439 | 0.65 |
|  | *Picea* | 25.9±8.75 | 39.3±23.2 | 13.6±4.72 | 1.186 | 0.323 |
|  | *V. vitis-idaea* | 4.52±2.64 | 6.39±2.26 | 7.68±1.94 | 1.842 | 0.178 |
|  | *V. myrtillus* | 5.94±3.71 | 14.4±6.04 | 16.2±5.06 | 3.106 | **0.062** |
|  | *Empetrum* | 4.72±2.07 | 6.62±2.69 | 11.0±6.57 | 0.541 | 0.588 |

**Supplementary Figures**

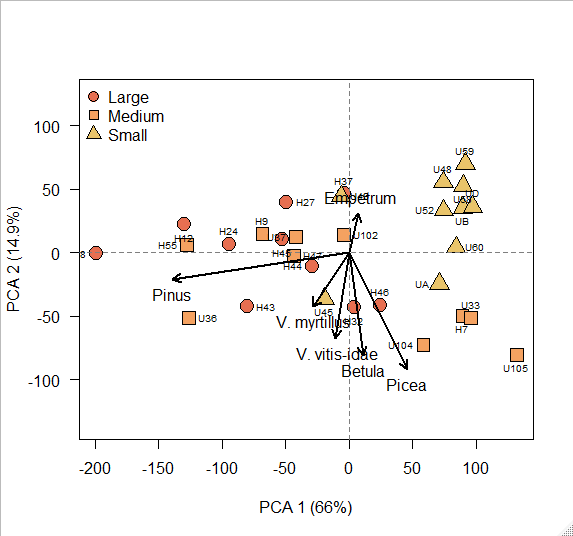

Fig S1: Results of a principal component analysis for the 30 islands (each represented as a separate data point), using the net primary production data (NPP) for the six dominant plant species in the system. Data on NPP are from Wardle et al., (2003, 2012). High values on the first PCA axis indicate that islands are dominated by resource conservative plant species such as *Picea* (loadings PC1: 0.3), while low values indicate that the community is dominated by more acquisitive species such as *Pinus* (loading PC1: -0.93; loading for the other species are Betula: 0.07; Empetrum: 0.05; V. vitis-idaea: -0.07; V. myrtillus: -0.19). Circles, squares and triangles indicate whether the island was in the large (>1.0 ha), medium (0.1 to 1.0 ha), or small (<0.1 ha size classes respectively).

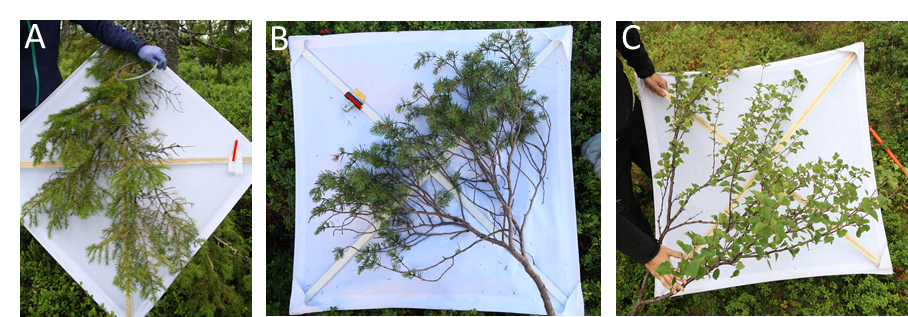

Fig. S2: Photographs of example branches of A) *Picea*, B) *Pinus*, and C) *Betula* on which invertebrates were sampled using the beat sheet method.

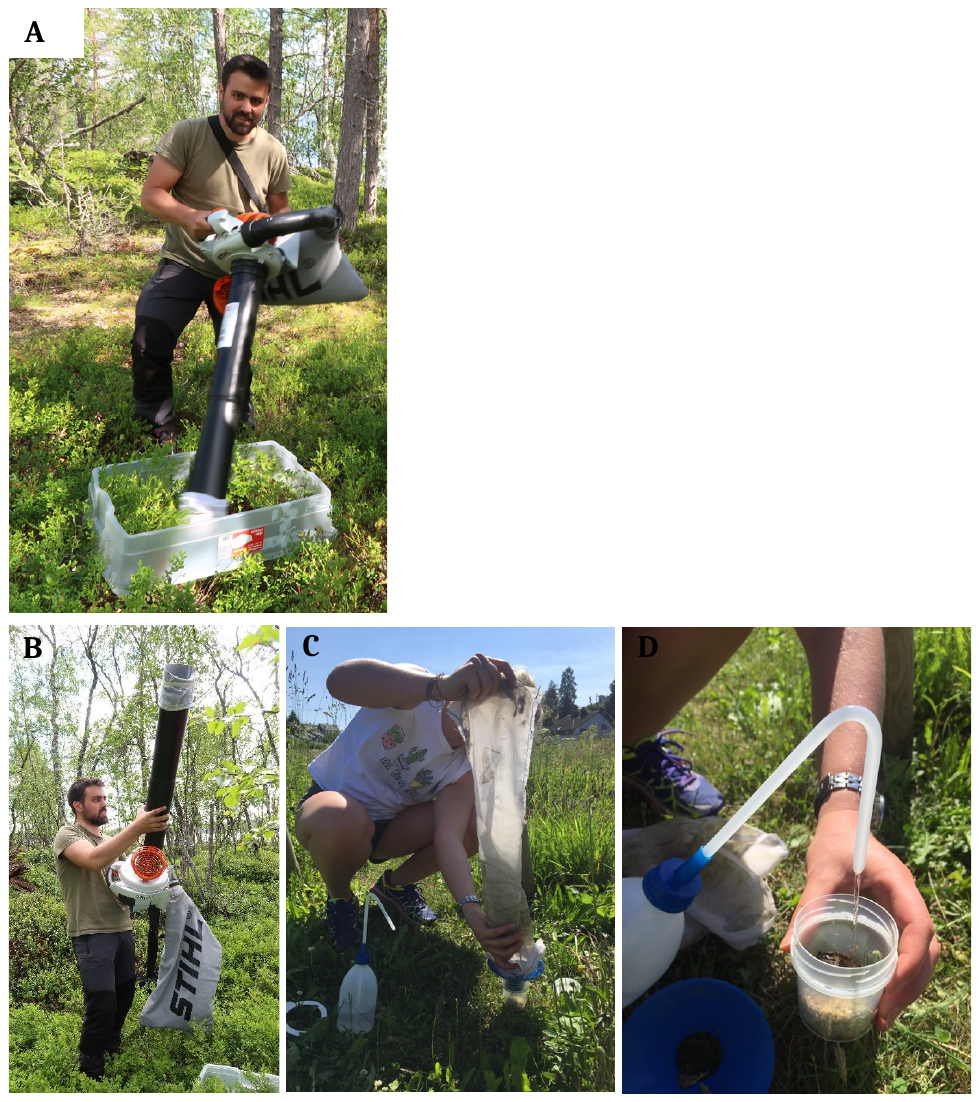

**Fig. S3.** A) Suction sampling was done for 100 s. B) After that, the nozzle was lifted upwards and the net removed. C) Invertebrates were transferred into a sampling container using a funnel with a wide opening. D) The sampling container was filled with 70 % ethanol. Pictures A and B by Anne Kempel, C and D by Tosca Mannall.

Fig. S4

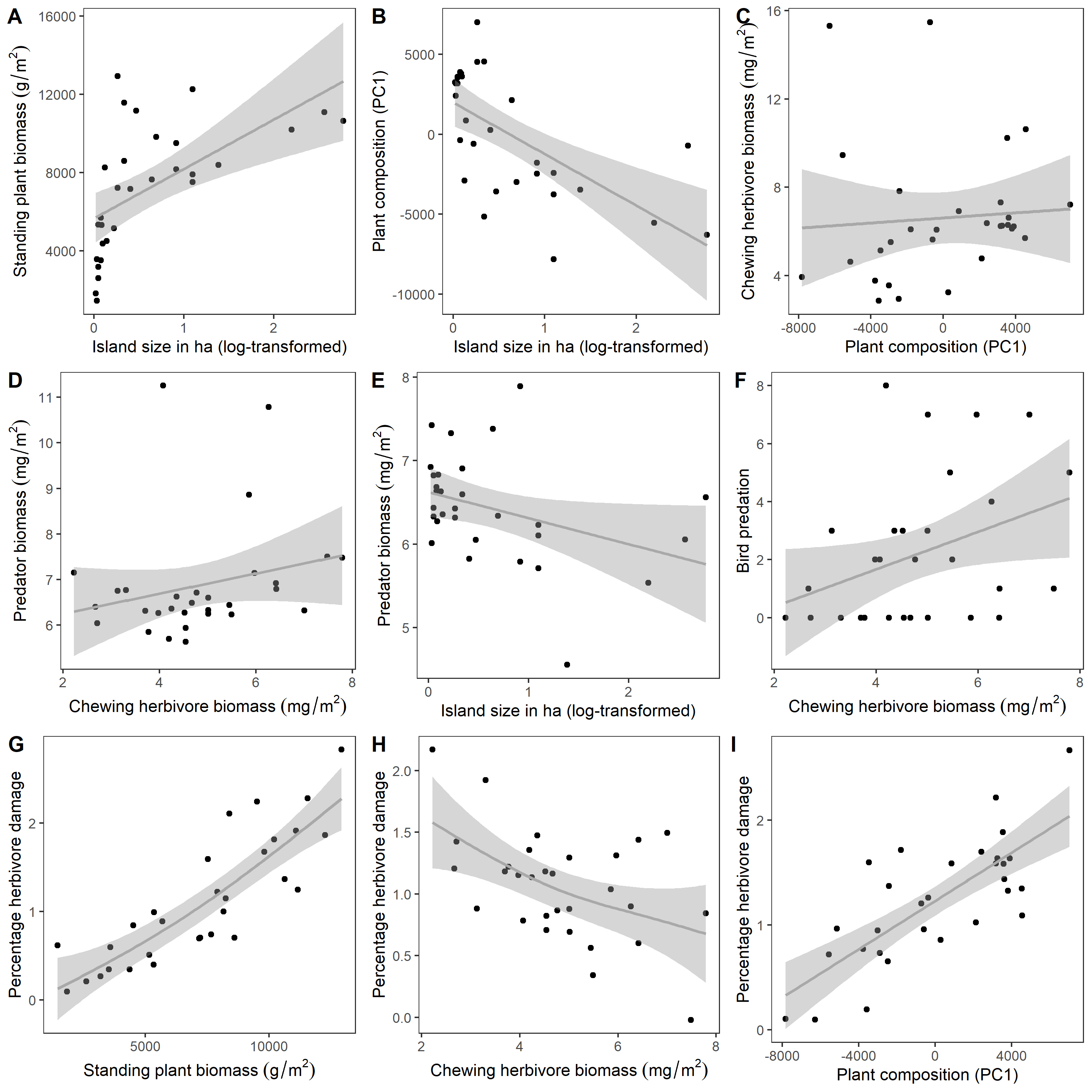

Fig. S4: Partial plot of the SEM in Fig. 4A. Impact of selected predictor variables on standing plant biomass (A), plant composition (B), chewing herbivore biomass (C), predator biomass (D,E), bird predation (F) and percentage herbivore damage on the phytometer seedlings (G-I), after removing all effects of all the other variables which are not plotted. Shaded areas represent 95% CI.

Fig. S5:

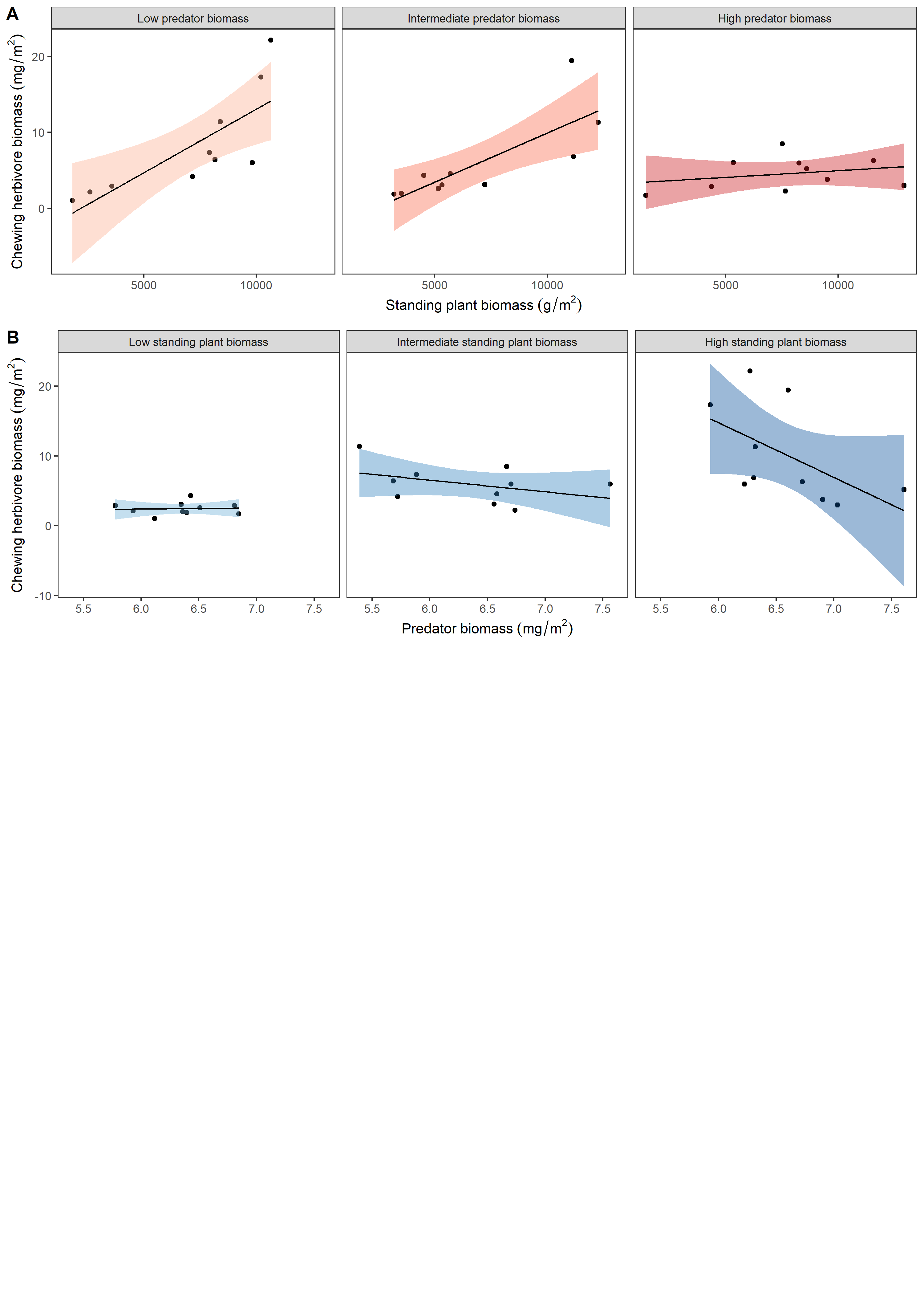

Fig. S5: Partial plots of the SEM in Fig. 4A, showing the interactive effect of standing plant biomass and predator biomass on chewing herbivore biomass, after removing all effects of all the other variables which are not plotted. A) Chewing herbivore biomass increases strongly with plant standing biomass for low predator biomass, but less strongly for high predator biomass. B) Chewing herbivore biomass is unresponsive to predator biomass for low standing plant biomass, but decreases for high plant biomass. For easier graphical visualisation, predator biomass (A) and plant standing biomass (B) has been binned into equal sized groups (low, medium and high), each group containing 10 islands. Shaded areas represent 95% CI.

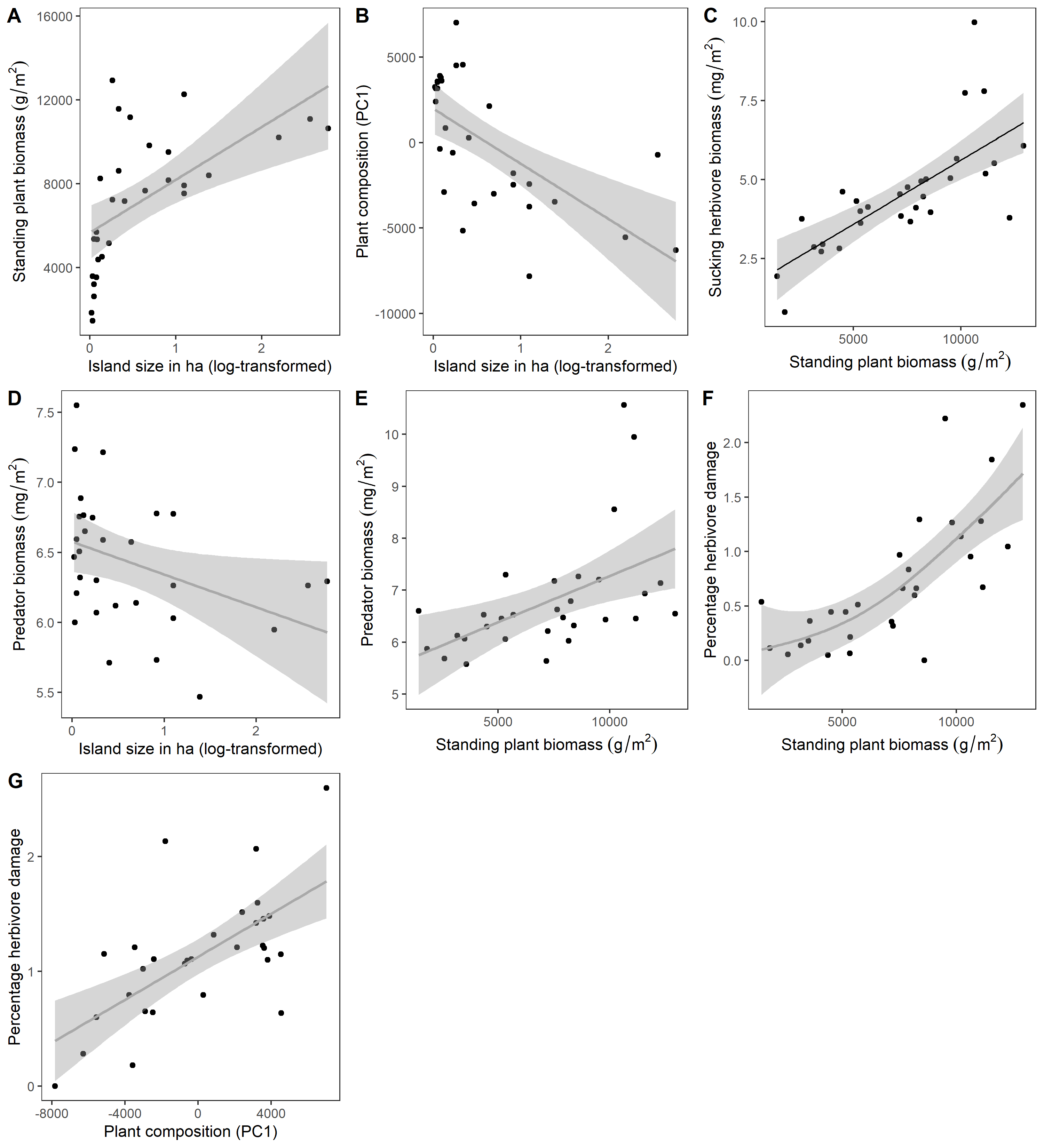

Fig. S6: Partial plot of the SEM in Fig. 4B. Impact of selected predictor variables (significant paths) on standing plant biomass (A), plant composition (B), sucking herbivore biomass (C), predator biomass (D,E) and percentage herbivore damage on the phytometer seedlings (F,G), after removing all effects of all the other variables which are not plotted. Shaded areas represent 95% CI.
